# FAP57/WDR65 targets assembly of a subset of inner arm dyneins and connects to regulatory hubs in cilia

**DOI:** 10.1101/688291

**Authors:** Jianfeng Lin, Thuc Vy Le, Katherine Augspurger, Douglas Tritschler, Raqual Bower, Gang Fu, Catherine Perrone, Eileen T. O’Toole, Kristyn VanderWaal Mills, Erin Dymek, Elizabeth Smith, Daniela Nicastro, Mary E. Porter

## Abstract

Ciliary motility depends on both the precise spatial organization of multiple dynein motors within the 96 nm axonemal repeat, and highly coordinated interactions between different dyneins and regulatory complexes located at the base of the radial spokes. Mutations in genes encoding cytoplasmic assembly factors, intraflagellar transport factors, docking proteins, dynein subunits, and associated regulatory proteins can all lead to defects in dynein assembly and ciliary motility. Significant progress has been made in the identification of dynein subunits and extrinsic factors required for pre-assembly of dynein complexes in the cytoplasm, but less is known about the docking factors that specify the unique binding sites for the different dynein isoforms on the surface of the doublet microtubules. We have used insertional mutagenesis to identify a new locus, *IDA8/BOP2*, required for targeting the assembly of a subset of inner dynein arms to a specific location in the 96 nm repeat. *IDA8* encodes FAP57/WDR65, a highly conserved WD repeat, coiled coil domain protein. Using high resolution proteomic and structural approaches, we find that FAP57 forms a discrete complex. Cryo-electron tomography coupled with epitope tagging and gold labeling reveal that FAP57 forms an extended structure that interconnects multiple inner dynein arms and regulatory complexes.

## Introduction

Cilia and flagella are microtubule-based organelles that play critical roles in cell motility and cell signaling, and defects in ciliary assembly, motility, or signaling can lead to a broad spectrum of diseases known as ciliopathies (reviewed in Reiter and Leroux, 2017). In vertebrates, ciliary motility is essential for the determination of the left-right body axis, development of the heart, movement of fluid in brain ventricles and spinal cord, clearance of mucus and debris in the respiratory tract, and sperm motility. Defects in motility can lead to situs inversus or heterotaxy, hydrocephalus and scoliosis, respiratory disease, and male infertility, symptoms often associated with primary ciliary dyskinesia (PCD) (Mitchison and Valente, 2017). Given the complexity of the microtubule-based 9+2 axonemal structure, motile ciliopathies are often under-diagnosed because of their genetic heterogeneity and multisystem variability (Werner et al., 2015). Yet many proteins of the ciliary axoneme are highly conserved (Li et al., 2004; Pazour et al., 2005; Albee et al., 2013), and so study of motile cilia in model organisms has provided insight into numerous genes and gene products potentially associated with PCD and other ciliopathies (Mitchison and Valente, 2017; Sigg et al., 2017).

Genomic and proteomic strategies have identified more than 600 proteins as structural components of the axoneme, and many other proteins contribute to the pre-assembly of axonemal complexes in the cytoplasm, their delivery to the basal body region, and their transport through the transition zone and into the ciliary compartment (reviewed in van Dam et al., 2019). Advances in high resolution imaging in combination with the ordered and repetitive nature of the axoneme structure have provided insight into the location of several axonemal complexes (Mizuno et al., 2012), but still only a third or so of the ciliary proteins have been clearly correlated with a specific structure.

Most motile cilia and flagella contain nine doublet microtubules (DMTs) that surround two central pair (CP) singlet MTs. The outer and inner dynein arms (ODA and IDA) are multi-subunit motors composed of heavy, intermediate and light chains (DHC, IC, LC) that form two distinct rows on the A-tubule of each DMT and generate the force for microtubule sliding (reviewed in King, 2018). The dynein motors are organized into a 96 nm functional unit that repeats along the length of the axoneme, with four ODAs and seven IDAs (I1/*f, a, b, c, e, g, d*) found at specific locations within each repeat. Dynein activity is coordinated by mechanical signals from the CP and its associated projections to a series of radial spokes that contact the DMTs near the base of the IDAs (Smith and Yang, 2004). The proximal to distal arrangement of the RS (RS1, RS2, RS3 or RS3S) and the multiple dyneins in each repeat is specified in part by two proteins (FAP59/FAP172 or CCDC39/CCDC40) that form a 96 nm ruler (Oda et al., 2014). The IC/LC complex of the I1/*f* dynein forms a regulatory node at the base of RS1, and the nexin-dynein regulatory complex (N-DRC) forms a second node at the base of RS2 (Gardner et al., 1994; Nicastro et al., 2006; Bower et al., 2009; Heuser et al., 2009; 2012) that is connected to the base of RS3/RS3S via the calmodulin and spoke-associated complex (CSC). The I1 dynein and N-DRC also connect to other structures in the 96 nm repeat and to the ODAs to coordinate dynein activity, but with the exception of the MIA complex next to the I1 dynein (Yamamoto et al., 2013), the identity of the connectors is largely unknown.

Here we identify a new group of mutations that alter ciliary motility and the assembly of a subset of IDAs in *Chlamydomonas*. Using plasmid rescue and a chromosome walk, we cloned and mapped the *IDA8* gene and found that it is linked to another motility mutation, *bop2-1*. We then characterized the molecular, biochemical, and structural phenotypes of the *ida8/bop2* mutations. We found that *IDA8* encodes FAP57, a highly conserved WD repeat and coiled coil protein also found in other species that have motile cilia with IDAs. Biochemical and proteomic analyses indicate that FAP57 is part of a sub-complex required for targeting or stabilizing the binding of a subset of IDAs. Thin section transmission electron microscopy (TEM) and cryo-electron tomography (cryo-ET) reveal the complexity of structural defects in *ida8/bop2* axonemes. Rescue with SNAP-tagged FAP57 constructs followed by streptavidin-gold labeling, sub-tomogram averaging, and image classification suggest that FAP57 forms an extended structure that interconnects multiple regulatory components and IDAs within the 96 nm axoneme repeat.

## Results

### Characterization of new ida mutations and identification of the IDA8/BOP2 locus

To identify novel genes required for assembly of the inner dynein arms (IDAs), we screened several collections of motility mutants in *Chlamydomonas* for strains that exhibited the slow swimming phenotype typical of *ida* mutants (Brokaw and Kamiya, 1991). Three strains characterized here, *ida8-1*, *ida8-2*, and *ida8-3*, were chosen for further study based on the similarities in their motility phenotypes and inner arm defects. All three strains swam forwards with an asymmetric waveform but their swimming velocities were reduced compared to wild-type strains (Supplemental Figure 1A), and their motility phenotypes co-segregated with their ability to grow on selective media. Thin section transmission electron microscopy (TEM) and 2D image averaging of isolated axonemes showed that structures in the IDA region were reduced (Supplemental Figure 1B). Genomic Southern blots indicated the presence of a single plasmid sequence in *ida8-1* and *ida8-3* (Supplemental Figure 1C).

To identify the gene that was disrupted by plasmid insertion, genomic DNA flanking the vector sequences was recovered by plasmid rescue (Materials and Methods). A unique 700 bp fragment designated flanking clone 1 (FC1) was recovered from *ida8-1* (Figure 1A). Southern blots of genomic DNA probed with FC1 confirmed the presence of a restriction fragment length polymorphism (RFLP) in *ida8-1*. FC1 was used to screen a phage library and isolate a series of overlapping clones spanning ∼40 kb of genomic DNA. Subclones were tested on Southern blots to determine the extent of DNA re-arrangement or deletion caused by the insertion events (Supplemental Figure 1D). The blots showed that the same genomic region was disrupted to varying degrees in all three *ida8* strains (Figure 1A). Transformation with a BAC clone (6h9) spanning this region rescued the motility defect (Supplemental Table 1).

**Figure 1.**
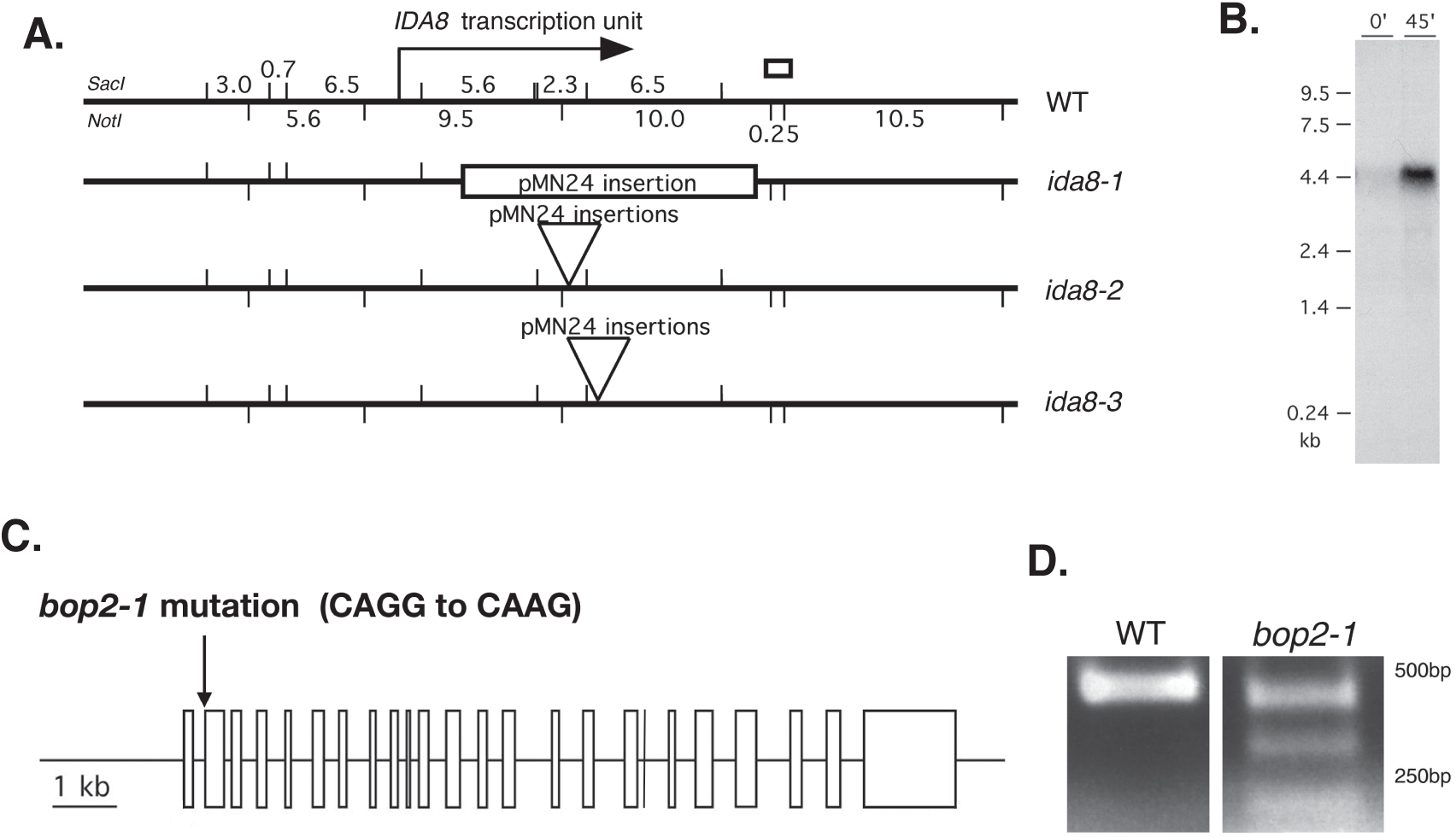
Molecular characterization of *ida8* and *bop2* mutations. **(A)** Diagram of the ∼40 kb region of genomic DNA around the *IDA8* locus in wild-type, with *SacI* restriction sites indicated on top and *Not1* restriction sites indicated below. The location of the *IDA8* transcription unit is shown by the arrow. The site of the genomic fragment recovered by plasmid rescue from *ida8-1* is shown by the white box. The next three lines show the sites of pMN24 insertion in each *ida8* allele as determined by Southern blotting (Supplemental Figure 1E). **(B)** Northern blot of total RNA isolated from WT cells before (0) and 45 minutes after deflagellation and probed with a 6.5 kb *SacI* restriction fragment that was missing in *ida8-1.* Other blots probed with the 5.6 kb and 2.3 kb *SacI* fragments and several RT-PCR products recognized the same transcript. **(C)** Diagram of the intron-exon structure of the *IDA8* gene showing the *bop2-1* mutation in the acceptor splice site of the second exon. **(D)** RT-PCR products obtained from wild-type and *bop2-1* RNA using primers surrounding the site of the *bop2-1* mutation were analyzed on an agarose gel. Sequence analysis identified premature stop codons in all of the RT-PCR products from *bop2-1*.

To determine the precise location of the *IDA8* transcription unit, subclones were used to probe Northern blots of wild-type RNA isolated before and after deflagellation. These blots defined an ∼12 kb region of genomic DNA that encodes an ∼5kb transcript whose expression was increased by deflagellation (Figure 1A, B). Transformation with a subclone containing the complete gene rescued the motility defects (Supplemental Table 1). DNA sequencing and RT-PCR revealed that the *IDA8* gene contains 24 exons (Figure 1C) that are predicted to encode a polypeptide of 1316 amino acid residues with an estimated molecular weight of ∼146 kD (Supplemental Figure 2A, B).

**Figure 2.**
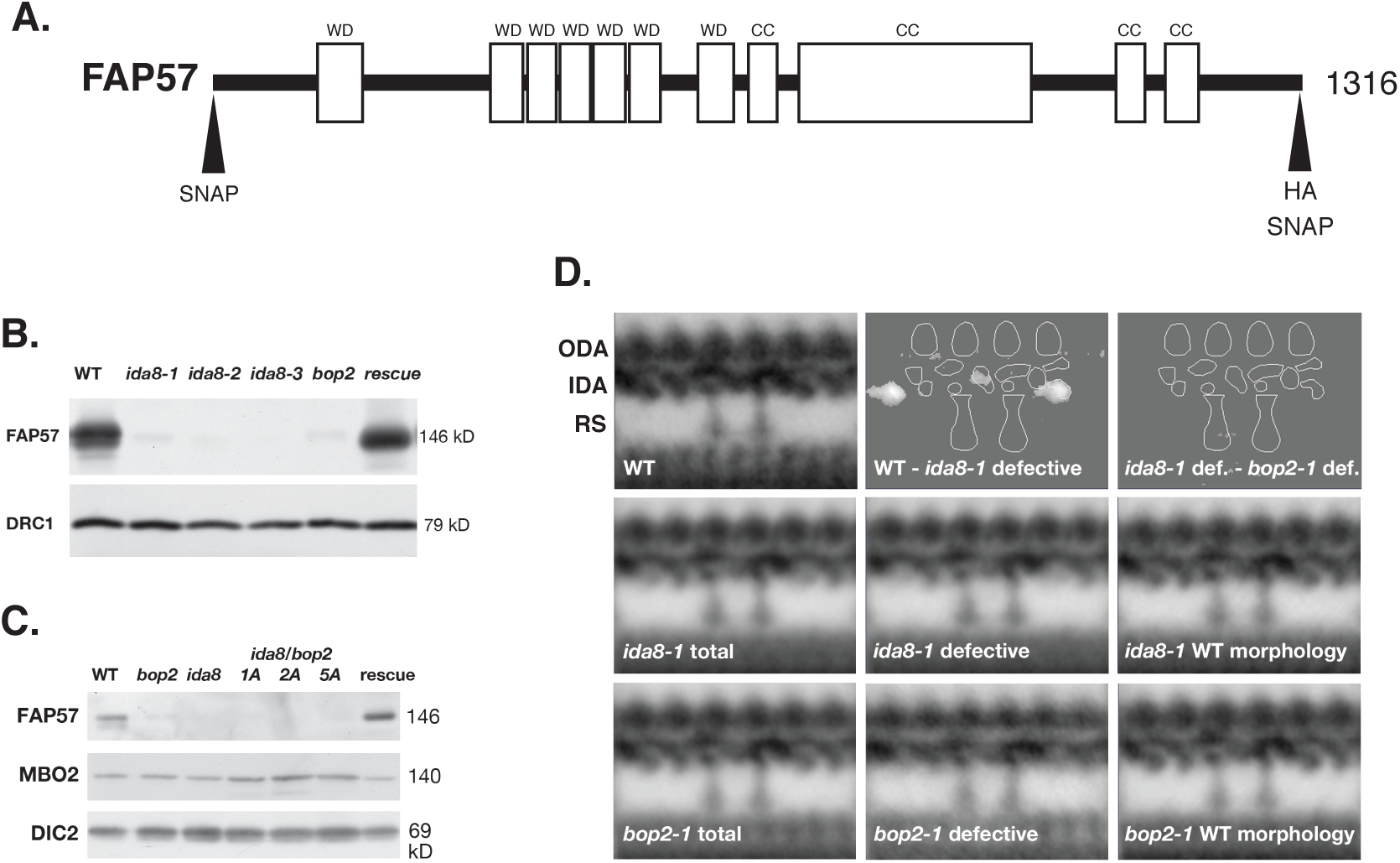
The *ida8* and *bop2* mutants have similar biochemical and structural defects. **(A)** Diagram of the *IDA8* gene product, FAP57, showing the location of the predicted WD repeat domains (WD) in the first half of the polypeptide and the coiled coil domains (CC) in the second half. Also shown are the positions of the HA and SNAP tags at the N-terminal and C-terminal ends. **(B)** A Western blot of axonemes from wild-type (WT), three *ida8* mutants, *bop2*-*1*, and an *ida8-1*; *FAP57* strain (rescue) was probed with antibodies against FAP57 and the N-DRC subunit DRC1 as a loading control. **(C)** A Western blot of axonemes from wild-type (WT), *bop2-1, ida8-1,* three *ida8-1/bop2-1* diploids (1A, 2A, 5A), and an *ida8-1*; *FAP57* strain (rescue) was probed with antibodies against FAP57, MBO2, and the outer arm DIC2 subunit, also known as IC69, as a loading control. **(D)** Averages and difference plots of the 96 nm repeat obtained by transmission electron microscopy (TEM) of thin sections and image averaging as described in O’Toole et al., 2005. The wild-type (WT) grand average (top left) was obtained from six axonemes with 65 repeats. The approximate locations of the outer dynein arms (ODA), inner dynein arms (IDA), and two radial spokes (RS) are indicated. The *ida8-1* grand average in the second row (*ida8-1* total) was obtained from 41 axonemes with 379 repeats. The *ida8-1* averages were divided into two classes, those with defective morphology (25 axonemes with 237 repeats) and those with WT morphology (16 axonemes with 142 repeats). The *bop2-1* grand average in the third row (*bop2-1* total) was obtained from 22 axonemes with 196 repeats. These were also separated into two classes, *bop2-1* defective (8 axonemes with 66 repeats) and *bop2-1* with WT morphology (14 axonemes with 130 repeats). The difference plots in the top row show identified two densities in the 96 nm repeat that were significantly different (P<0.05) between the WT and *ida8-1* defective grand averages. No significant difference was detected between the *ida8-1* defective average and *bop2-1* defective average.

Genetic and molecular mapping further revealed that the *IDA8* locus was located on the left arm of Chromosome 4 (Materials and Methods), close to the motility mutation *bop2-1* (Dutcher et al., 1988). Sequencing of *bop2-1* DNA identified a single base pair mutation at nucleotide #605 (CAGG to CAAG) in the acceptor splice site of exon 2 (Figure 1C). Three alternatively spliced transcripts were detected by RT-PCR (Figure 1D), and sequence analysis revealed that all three transcripts contained frameshifts that resulted in premature stop codons. Diploid strains containing *bop2-1* and *ida8-1* displayed the same motility phenotype as the parental strains, and transformation of *bop2-1* with the p59c2 subclone rescued the motility defects (Supplemental Table 1).

### The IDA8/BOP2 locus encodes the conserved WD repeat and coiled-coil containing polypeptide FAP57

The *IDA8/BOP2* gene encodes a polypeptide identified in the flagellar proteome as FAP57 (Pazour et al., 2005). FAP57 is predicted to contain an N-terminal region with several WD repeat domains (residues 1-622) and C-terminal region with several coiled-coil domains (residues 640-1188) and more variable, low complexity domains (residues 1206-1314) that are potentially disordered (Figure 2A). Both the polypeptide sequence and structural domains are highly conserved in other species with motile cilia and flagella (Nevers et al., 2017), including a polypeptide identified as WDR65 in vertebrates, with whom it shares almost 42% sequence identity and 61% sequence similarity. Interestingly, FAP57 orthologues are found in organisms that only assemble IDAs, such as *Physcomitrella* (Table 1), but not in species that only assemble ODAs, such as *Thalissiosira*.

**Table 1.**
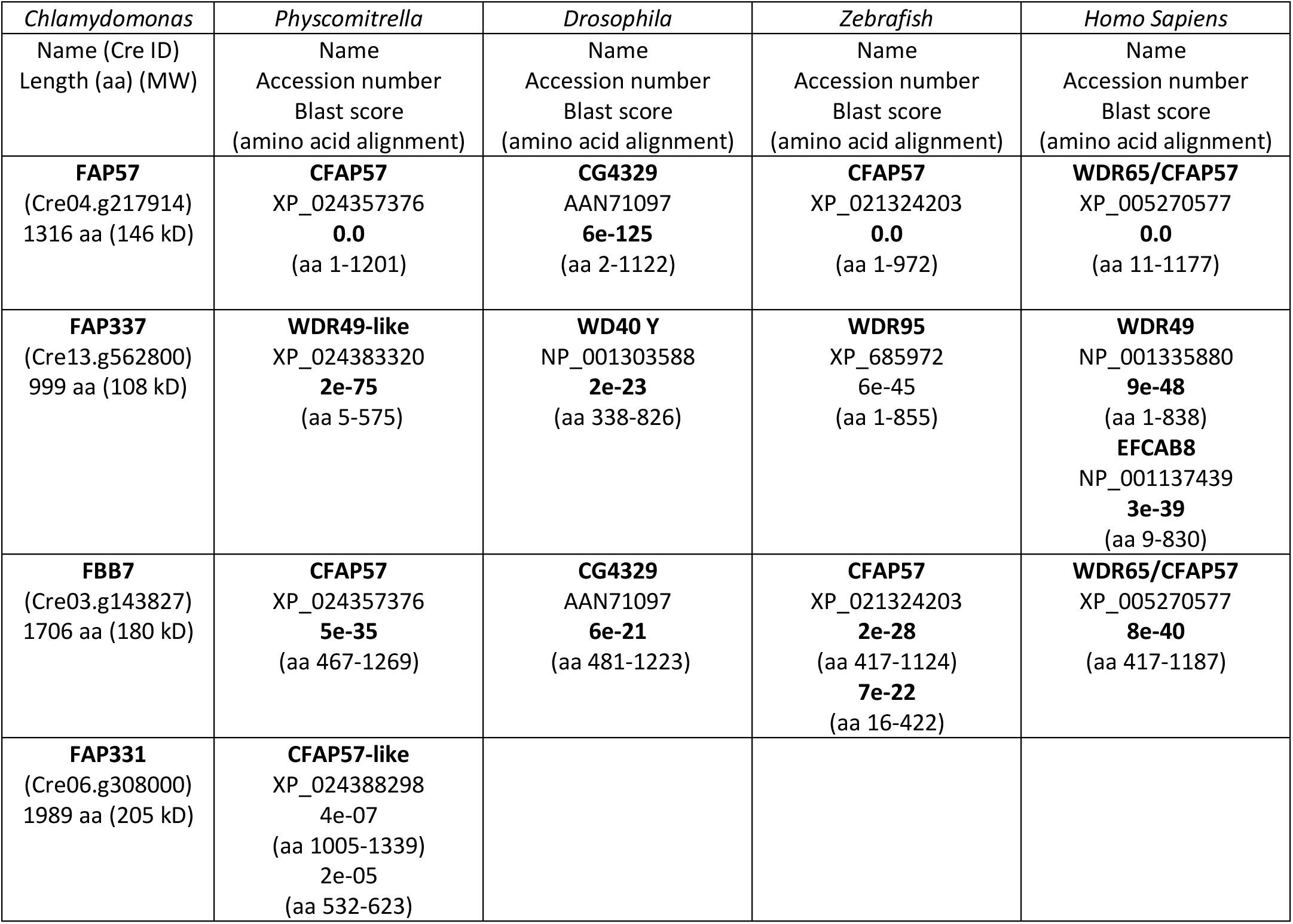
Orthologues of polypeptides altered in *ida8*.

| *Chlamydomonas* | *Physcomitrella* | *Drosophila* | *Zebrafish* | *Homo Sapiens* |
| --- | --- | --- | --- | --- |
| Name (Cre ID)<br>Length (aa) (MW) | Name<br>Accession number<br>Blast score<br>(amino acid alignment) | Name<br>Accession number<br>Blast score<br>(amino acid alignment) | Name<br>Accession number<br>Blast score<br>(amino acid alignment) | Name<br>Accession number<br>Blast score<br>(amino acid alignment) |
| **FAP57**<br>(Cre04.g217914)<br>1316 aa (146 kD) | **CFAP57**<br>XP_024357376<br>**0.0**<br>(aa 1-1201) | **CG4329**<br>AAN71097<br>**6e-125**<br>(aa 2-1122) | **CFAP57**<br>XP_021324203<br>**0.0**<br>(aa 1-972) | **WDR65/CFAP57**<br>XP_005270577<br>**0.0**<br>(aa 11-1177) |
| **FAP337**<br>(Cre13.g562800)<br>999 aa (108 kD) | **WDR49-like**<br>XP_024383320<br>**2e-75**<br>(aa 5-575) | **WD40 Y**<br>NP_001303588<br>**2e-23**<br>(aa 338-826) | **WDR95**<br>XP_685972<br>6e-45<br>(aa 1-855) | **WDR49**<br>NP_001335880<br>**9e-48**<br>(aa 1-838)<br>**EFCAB8**<br>NP_001137439<br>**3e-39**<br>(aa 9-830) |
| **FBB7**<br>(Cre03.g143827)<br>1706 aa (180 kD) | **CFAP57**<br>XP_024357376<br>**5e-35**<br>(aa 467-1269) | **CG4329**<br>AAN71097<br>**6e-21**<br>(aa 481-1223) | **CFAP57**<br>XP_021324203<br>**2e-28**<br>(aa 417-1124)<br>**7e-22**<br>(aa 16-422) | **WDR65/CFAP57**<br>XP_005270577<br>**8e-40**<br>(aa 417-1187) |
| **FAP331**<br>(Cre06.g308000)<br>1989 aa (205 kD) | **CFAP57-like**<br>XP_024388298<br>4e-07<br>(aa 1005-1339)<br>2e-05<br>(aa 532-623) |  |  |  |

To assess the location and distribution of the FAP57 polypeptide in wild-type and mutant *Chlamydomonas* strains, we generated a specific antibody against a conserved peptide sequence at amino acid residues 460-480. Western blots showed that the affinity purified antibody detected a band of ∼146 kD present in wild-type and rescued strains but missing in axonemes from *ida8*, *bop2-1,* and *ida8-1/bop2-1* diploids (Figures 2B, C). FAP57 therefore corresponds to the ∼152 kD band previously described as missing on gels of *bop2-1* axonemes (King et al., 1994).

Because previous study of *bop2-1* had indicated radial asymmetry in the assembly of uncharacterized inner arm structures (King et al., 1994), we directly compared the defects in *ida8-1* and *bop2-1* axonemes by thin section TEM of longitudinal sections and 2D averaging of the 96 nm repeat. As shown in Figure 2D, both mutants showed heterogeneity in the assembly of structures in the inner arm region. When sorted into two classes, a significant number of *ida8-1* (37%) and *bop2-1* (66%) repeats were similar to wild type, but several *ida8-1* (63%) and *bop2-1* (34%) repeats were also missing two structures, one close to the base of the I1/*f* dynein between RS1 and RS2, and a second at the distal end of the 96 nm repeat.

### FAP57 is located in the basal body region and along the length of the axoneme

To better understand the role of FAP57 in the assembly of inner dynein arm structures, we generated epitope-tagged constructs of FAP57 (Supplemental Figure 2C, D) and used these constructs and the affinity-purified FAP57 antibody to analyze the distribution of FAP57 in wild-type, mutant, and rescued cells. Transformation of *ida8-1* and *bop2-1* with a *FAP57-HA* construct (Figure 2A) restored near wild-type motility in both strains (Figure 3A, Supplemental Movies 1-5). The HA-tagged protein assembled into axonemes and migrated at ∼151 kD on Western blots (Figure 3B). Localization by immunofluorescence microscopy of fixed cells revealed that FAP57-HA was concentrated near the basal bodies but also found along the entire length of the axoneme (Figure 3C). The intense staining of FAP57-HA at the basal body region was qualitatively different from that observed with another HA-tagged axonemal polypeptide, such as DRC4-HA, one of the subunits of the N-DRC (Figure 3C). Staining of the isolated nuclear-flagellar apparatus indicated that FAP57 was stably associated with isolated basal bodies and axonemes, but not present in the transition zone (Figure 3D).

**Figure 3.**
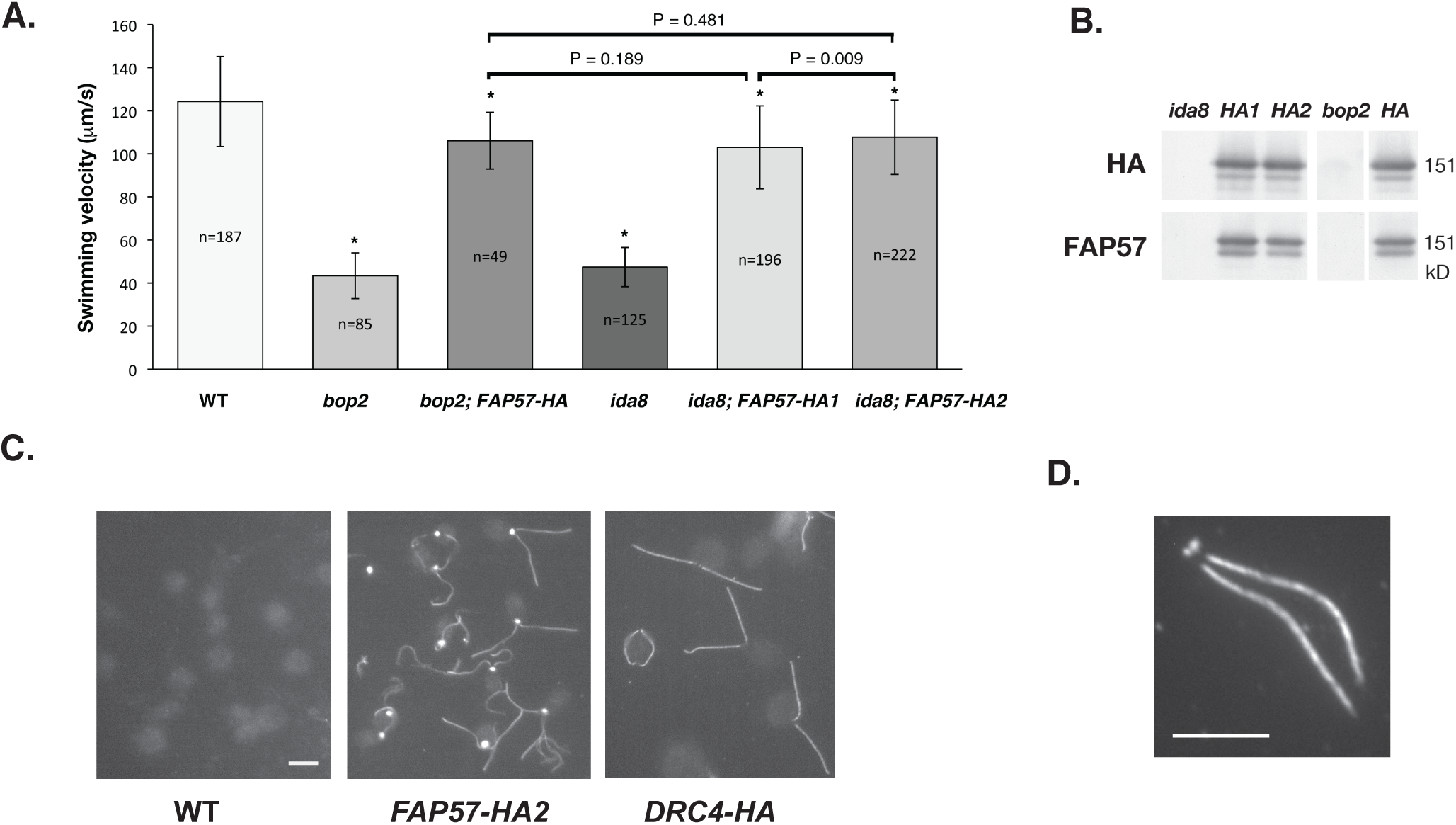
Rescue of *bop2-1* and *ida8-1* with FAP57-HA reveals its subcellular location. **(A)** Measurements of forward swimming velocity demonstrated that the speed of *bop2-1* and *ida8-1* transformants increased to near wild-type levels following rescue with the HA-tagged *FAP57* gene. **(B)** A Western blot of axonemes from *ida8-1*, two *ida8-1; FAP57-HA* rescued strains (*HA1* and *HA2*), *bop2-1*, and a *bop2-1; FAP57-HA* rescued strain (*HA*) was probed with antibodies against the HA tag and FAP57. **(C)** Immunofluorescence images of fixed cells stained with an HA antibody revealed background staining of cell bodies in wild-type (WT) and bright staining of the basal body region and two flagella in *ida8-1; FAP57-HA* rescued cells. Images of *pf2-4; DRC4-HA* rescued cells showed antibody staining of the two flagella but much weaker staining of the basal body region. (Scale bar = 5 μm). **(D)** An immunofluorescence image of a nuclear flagellar apparatus (NFAP) obtained by autolysin treatment and detergent extraction of an *ida8-1; FAP57-HA* rescued strain. (Scale bar = 5 μm).

### Biochemical fractionation of FAP57 suggests a specific association with a subset of IDA isoforms

To determine if FAP57 might be associated with a specific sub-complex of axonemal polypeptides, we probed blots of axonemes isolated from several classes of motility mutants. These included outer arm mutants (*pf22, pf28, sup-pf2*), inner arm mutants (*pf23, pf9, ida4, mia1, mia2*), central pair mutants (*pf19, pf6*), nexin-dynein regulatory complex (N-DRC) mutants (*pf2, pf3, sup-pf3, sup-pf4*), and other motility mutants with more symmetric waveforms (*mbo1, pf12*). As shown in Supplemental Figure 3A-C, FAP57 was detected at near wild-type levels in all of these strains.

Because *bop2-1* was originally isolated as an extragenic suppressor of *pf10* (Dutcher et al., 1998), we also analyzed the phenotypes of the double mutants *bop2-1; pf10* and *ida8-1; pf10*. Western blots confirmed that FAP57 was present in *pf10* axonemes but missing in the double mutants (Supplemental Figure 4A). High speed movies showed that *pf10* cells swam in small circles with a more symmetric waveform than the asymmetric breast stroke typically executed by wild-type, *ida8*, and *bop2* cells. Double mutant cells *bop2-1; pf10* and *ida8-1; pf10* swam with slightly more asymmetric waveforms than *pf10*, but still did not make significant forward progress (Supplemental Figure 4B, Supplemental Movies 6-8). As an alternative test for dynein activity, we measured DMT sliding velocities using protease-treated axonemes and an *in vitro* sliding disintegration assay. As shown in Supplemental Figure 4C, the microtubule sliding velocities of *bop2* and *ida8-1* axonemes were significantly slower than WT axonemes. Likewise, the sliding velocities of the *ida8-1; pf10* and *bop2; pf10* double mutants were slower than *pf10*. These results, taken together with previous epistasis tests indicating that the *bop2-1* mutation enhanced the motility defects observed with other dynein or *n-drc* mutants (King et al., 1994), suggested that FAP57 was part of a previously uncharacterized axonemal sub-complex involved in the assembly, transport, targeting, and/or regulation of IDAs.

To characterize the biochemical properties of FAP57, we subjected wild-type flagella to a series of extraction protocols and analyzed the resulting extracts on silver stained gels and/or Western blots. As shown in the Western blot in Supplemental Figure 3D, FAP57 was readily detected in isolated flagella, but not significantly extracted with either non-ionic detergents, which solubilizes membrane plus matrix proteins, or with 10 mM MgATP, which typically extracts IFT motor proteins (Cole et al., 1998; Pazour et al., 1999; Perrone et al., 2003). However, extraction of axonemes with 0.6M NaCl followed by 0.5M NaI solubilized nearly all of the FAP57 (Supplemental Figure 3D). Sequential treatment of axonemes with 0.2M, 0.4M, and 0.6M NaI showed that FAP57 was more resistant to extraction than the I1 dynein subunit IC140, but more readily solubilized than the radial spoke subunit RSP16 (Figure 4A). Fractionation of dynein extracts by sucrose density gradient centrifugation indicated that FAP57 sedimented at ∼8-10S, which is slower than either I1 dynein (IC140) at ∼20S or dynein *c* (DHC9) at ∼12-13S (Figure 4B). Fractionation of *pf28* extracts by FPLC chromatography revealed that FAP57 eluted with a broad profile, co-fractionating with IDAs in peaks *d, e, f,* and *g* (Figure 4C). The highest concentration of FAP57 coincided with the peak of dynein *g*. The elution profile of FAP57 was unchanged in dynein extracts obtained from *pf10, mbo1,* and *ida4* axonemes (data not shown). Because FAP57 was not reduced in mutants that lack dynein *d, e,* or *f* (i.e., *ida4, pf3, pf9,* see Supplemental Figure 3A, B), it seemed likely that FAP57 is not a *bona fide* dynein subunit but may be peripherally associated with the IDAs as a docking factor.

**Figure 4.**
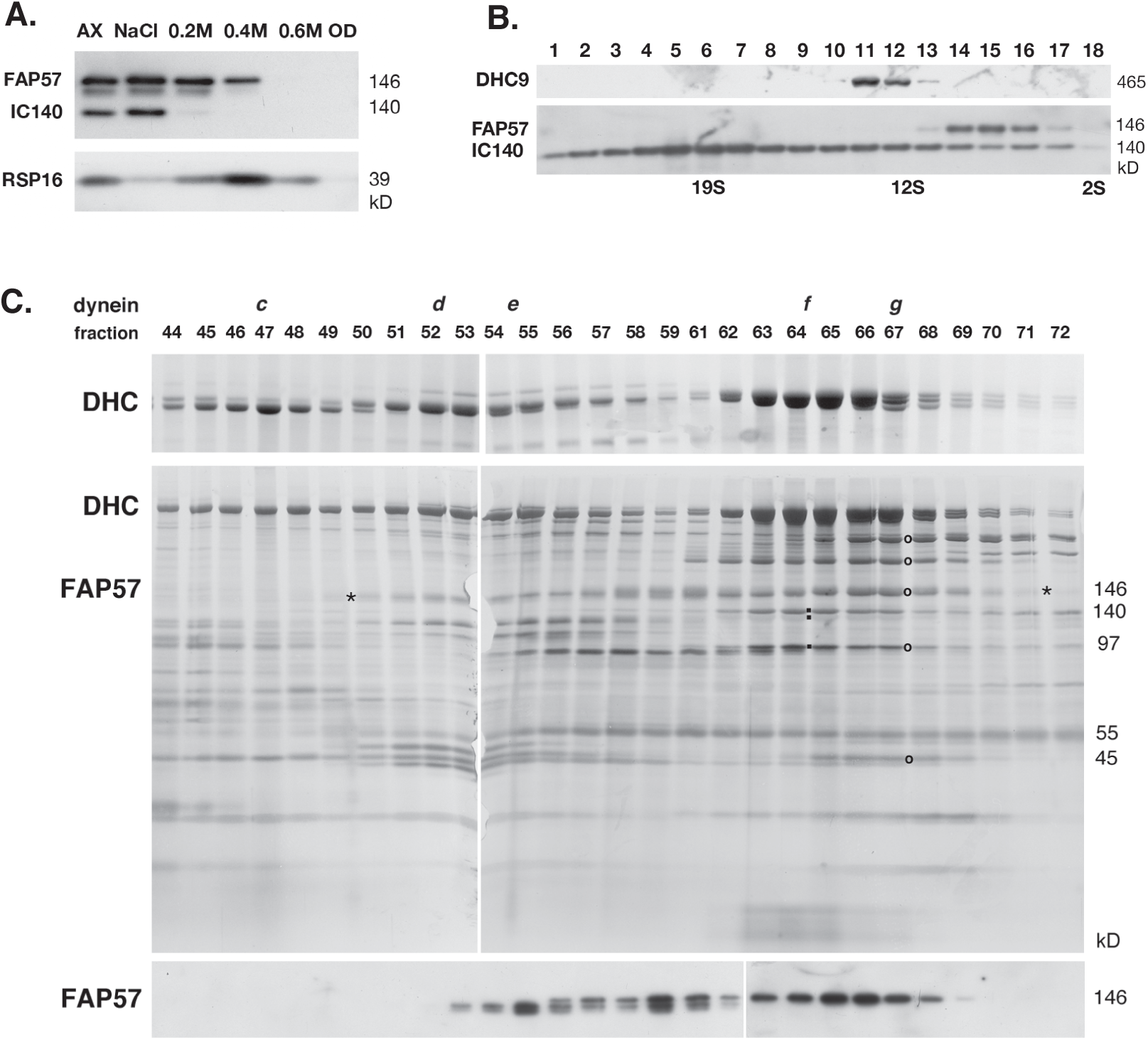
Biochemical fractionation of FAP57 demonstrates co-extraction and co-elution with a subset of IDAs. **(A)** A Western blot of wild-type axonemes (AX) and extracts obtained by sequential treatment with 0.6M NaCl, 0.2M, 0.4M, and 0.6M NaI, and the final pellet of extracted outer doublets (OD) was probed with antibodies against FAP57, the I1 dynein subunit IC140, and the radial spoke subunit RSP16. **(B)** A dynein extract was fractionated by sucrose density gradient centrifugation. Fractions 1-18 were analyzed on a Western blot probed with antibodies against DHC9, FAP57, and IC140. **(C)** A dynein extract from the outer arm mutant p*f28* was fractionated by FPLC chromatography. Fractions 44-72 were analyzed by SDS-PAGE on 3-5% (top) and 5-15% (middle) gels stained with silver or on a Western blot (bottom) stained with the FAP57 antibody. FAP57 eluted in a broad region (see asterisks) that overlapped with the FPLC peaks of dyneins *d*, *e*, *f*, and *g*. The small black squares indicate the dynein ICs associated with I1/*f* dynein. The black circles indicate the bands in peak *g* that were analyzed by MS/MS.

### Identification of polypeptides associated with FAP57 by mass spectrometry

To identify other polypeptides that might interact with FAP57, we analyzed FPLC fractions and wild-type and mutant axonemes by mass spectrometry (MS/MS). For the FPLC samples, fractions spanning the peaks of dynein *f* and *g* were separated by SDS-PAGE and silver stained (Figure 4C). Prominent bands were excised from the gel, digested with trypsin, and analyzed by MS/MS. As expected, numerous peptides from FAP57 and co-purifying dynein subunits were readily detected. Five other polypeptides co-eluted with FAP57; these included FAP44, FAP43, FAP244, FAP159, and FAP75, in order of peptide abundance (Supplemental Table 4). Little is known about FAP159 and FAP75, but FAP43, FAP44, and FAP244 have been identified as subunits of a tether-tether head (T/TH) complex linking the I1 dynein motor domains to the DMT and also interacting with the base of dynein *d* (Fu et al., 2018; Kubo et al., 2018; Urbanska et al., 2018). FAP43, FAP44, and FAP244 share structural similarity with FAP57 with respect to the arrangement of their WD repeat and coiled coil domains (Fu et al., 2018; Kubo et al., 2018), and FAP43 and FAP44 have been proposed to interact with FAP57 in *Tetrahymena* cilia based on proximity labeling (Urbanska et al., 2018).

To see if any of the polypeptides that co-eluted with FAP57 might be altered in *ida8*, axonemes from *ida8-1* and an HA-rescued strain were labeled in duplicate using four different iTRAQ tags, digested, fractionated by liquid chromatography, and analyzed by MS/MS to identify the total complement of polypeptides present in each sample (see Materials and Methods). The protein ratios were then analyzed to identify those polypeptides whose *ida8/HA* ratios were significantly different (P < 0.05) from the control ratio (*HA/HA*). Several proteins were reduced to variable degrees, but only two polypeptides, FAP57 and Cre13.g562800, were reproducibly and significantly reduced below 30% in *ida8-1* axonemes (Table 2). Cre13.g562800, is an EF hand, WD repeat containing polypeptide identified in the flagellar proteome as FAP337 (Pazour et al., 2005). It is closely related to another protein in *Chlamydomonas*, Cre07.g313850 (Blast score 1e-50), and also shares significant sequence homology with two vertebrate proteins WDR49 and EFCAB8 (Table 1). The iTRAQ ratios of two other proteins were increased more than 50% in *ida8-1*, FBB7 (Cre03.g143827) and FAP331 (Cre06.g308000). Both contain N-terminal regions with multiple WD repeats and C-terminal regions with coiled coil domains, similar to FAP57. To verify the changes in protein composition predicted by iTRAQ analysis, we also fractionated axonemes from WT, *ida8*, and the HA rescued strain by SDS-PAGE, cut bands from the appropriate regions, digested the samples with trypsin, and analyzed both the number of unique peptides and total spectra using label free quantification. Spectral counting confirmed that FAP57 and FAP337 were significantly reduced in *ida8* (<10% of WT), that FAP331 and FBB7 were increased in *ida8* (>30% of WT), and that all were restored to WT levels in *FAP57-HA* rescued axonemes. The organization of polypeptide domains in each of these proteins is shown diagrammatically in Supplemental Figure 5.

**Table 2.** iTRAQ protein ratios in *ida8* and *FAP57-HA* axonemes.

| Protein | Experiment 1 |  |  | Experiment 2 |  |  |
| --- | --- | --- | --- | --- | --- | --- |
|  | Peptides* | HA/HA | *ida8*/HA | Peptides* | HA/HA | *ida8*/HA |
| Reduced (<0.30) |  |  |  |  |  |  |
| FAP57 | 76 (94) | 0.93 | **0.11** | 100 (172) | 0.89 | **0.16** |
| Cre13.g562800 (FAP337) | 11 (13) | 1.00 | **0.26** | 17 (21) | 0.98 | **0.26** |
| Increased (>1.50) |  |  |  |  |  |  |
| Cre06.g308000 (FAP331) | 55 (58) | 1.12 | **2.01** | 55 (67) | 1.01 | **1.59** |
| Cre03.g143827 (FBB7) | 59 (71) | 1.16 | **2.45** | 56 (69) | 1.03 | **1.83** |
| **Dynein heavy chains** |  |  |  |  |  |  |
| 1-alpha DHC | 318 (368) | 1.01 | 1.02 | 357 (660) | 0.98 | 0.89 |
| 1-beta DHC | 328 (419) | 1.02 | 1.00 | 368 (735) | 0.97 | 0.87 |
| DHC2 | 201 (238) | 0.99 | **0.86** | 286 (508) | 0.96 | **0.66** |
| DHC3 | 76 (69) | 0.98 | **0.54** | 99 (113) | 0.95 | **0.45** |
| DHC4 | 144 (122) | 1.02 | 1.16 | 160 (167) | 0.97 | 0.84 |
| DHC5 | 212 (216) | 1.03 | 1.15 | 240 (355) | 0.96 | 0.91 |
| DHC6 | 182 (202) | 1.02 | 1.07 | 237 (395) | 0.97 | 0.77 |
| DHC7 | 209 (235) | 0.99 | **0.63** | 259 (426) | 0.95 | **0.40** |
| DHC8 | 216 (229) | 1.02 | 1.08 | 251 (390) | 0.97 | 0.86 |
| DHC9 | 268 (299) | 1.01 | 1.08 | 290 (355) | 0.98 | 0.91 |
| DHC11 | 146 (135) | 1.03 | 1.05 | 173 (213) | 0.97 | 0.89 |
| DHC12 | 78 (96) | 1.02 | 1.03 | 100 (129) | 0.99 | 0.83 |
| Alpha DHC | 505 (516) | 1.02 | 1.10 | 531 (1084) | 0.98 | 0.88 |
| Beta DHC | 558 (625) | 1.02 | 1.12 | 605 (1321) | 0.97 | 0.89 |
| Gamma DHC | 375 (428) | 1.02 | 1.13 | 446 (907) | 0.98 | 0.89 |
| **I1 dynein associated** |  |  |  |  |  |  |
| IC140/DIC3 | 68 (119) | 1.01 | 1.04 | 55 (79) | 1.01 | 1.10 |
| IC138/DIC4 | 76 (127) | 0.96 | 1.01 | 64 (77) | 1.04 | 1.29 |
| IC97 | 56 (118) | 0.99 | 1.04 | 41 (62) | 1.00 | 1.15 |
| MIA1/FAP100 | 35 (35) | 1.01 | 1.06 | 42 (53) | 0.99 | 0.93 |
| MIA2/FAP73 | 23 (26) | 1.01 | 0.98 | 23 (38) | 1.01 | 0.99 |
| FAP43 | 87 (107) | 1.03 | 1.06 | 95 (184) | 0.98 | 0.94 |
| FAP44 | 104 (117) | 1.05 | 1.10 | 117 (212) | 0.97 | 0.91 |
| FAP244 | 23 (24) | 0.97 | 0.99 | 80 (122) | 0.97 | 0.86 |
| **N-DRC** |  |  |  |  |  |  |
| DRC2 | 36 (54) | 1.00 | 1.03 | 36 (68) | 1.02 | 0.88 |
| DRC3 | 47 (59) | 1.03 | 1.03 | 62 (118) | 0.99 | 0.90 |
| DRC4 | 65 (79) | 1.00 | 0.99 | 73 (138) | 0.99 | 1.03 |
| DRC5 | 21 (18) | 0.96 | 0.96 | 31 (27) | 0.94 | 0.96 |
| DRC6 | 23 (34) | 0.97 | 0.99 | 21 (42) | 0.99 | 0.94 |
| DRC7 | 120 (172) | 1.02 | 1.05 | 133 (304) | 0.98 | 0.92 |
| DRC8 | 17 (25) | 1.05 | 1.00 | 13 (30) | 0.97 | 1.00 |
| DRC9 | 33 (39) | 1.03 | 1.07 | 35 (69) | 1.02 | 0.99 |
| DRC10 | 24 (37) | 0.98 | 0.98 | 29 (56) | 1.00 | 0.98 |
| DRC11 | 72 (93) | 1.00 | 0.95 | 71 (136) | 0.98 | 0.96 |
| **MT doublet associated** |  |  |  |  |  |  |
| Rib43 | 52 (52) | 0.95 | 0.86 | 42 (88) | 1.03 | 1.08 |
| Rib72 | 159 (181) | 1.02 | 0.96 | 174 (333) | 0.97 | 0.95 |
| FAP59 | 62 (68) | 1.04 | 0.99 | 61 (117) | 0.96 | 0.90 |
| FAP172 | 67 (71) | 1.01 | 0.95 | 61 (113) | 0.97 | 0.84 |
| MBO2 | 74 (91) | 1.03 | 1.02 | 81 (139) | 0.98 | 0.90 |
| TUA | 449 (440) | 0.98 | 0.98 | 569 (1202) | 1.00 | 1.00 |
| TUB | 584 (632) | 1.02 | 1.03 | 723 (1160) | 0.99 | 1.01 |
| FAP20 | 49 (89) | 1.05 | 1.12 | 37 (36) | 1.01 | 1.12 |
| PACRG | 76 (268) | 0.98 | 1.12 | 68 (95) | 1.01 | 1.13 |

\*Peptides identified at the 95% confidence interval (peptides used for quantification). The HA/HA ratio represents variation between technical replicates of the same biological sample (typically less than 10%). The *ida8*/HA ratio is an average of two technical replicates of *ida8* relative to the *FAP57-HA* rescued strain. The two experiments are different biological replicates. The *ida8*/HA ratios that were significantly different ( $P < 0.05$ ) in both experiments are highlighted in bold.

Because *ida8-1* and *bop2-1* displayed defects in the assembly of IDA structures located in the 96 nm repeat, we also analyzed the iTRAQ ratios of several proteins previously localized near the IDAs (Table 2). All polypeptides associated with the two-headed I1/*f* dynein, which is located at the proximal end of the 96-nm repeat (Piperno et al., 1990; Mastronarde et al., 1992), were present at wild-type levels. These include the 1*α* and 1*β* DHCs, several I1 intermediate chains, the two MIA proteins (FAP73 and FAP100) implicated in the regulation of I1 dynein (Yamamoto et al., 2013), and the three T/TH proteins, FAP43, FAP44, and FAP244, mentioned above. All subunits of the N-DRC, which is located at the distal end of the repeat (Gardner et al., 1994; Heuser et al., 2009), were also present at wild-type levels. Moreover, the ratios of several coiled-coil proteins were also unchanged in *ida8-1*. These include Rib43a and Rib72, two proteins found inside the lumen of the A-tubule (Stoddard et al., 2018) and the 96nm ruler complex FAP59/FAP172, which has been identified as elongated structure that is tightly associated with the DMTs, establishes the dimensions of the 96 nm repeat, and helps to specify the binding sites of the radial spokes and IDAs (Oda et al., 2014). No significant changes were observed in any of the outer arm DHCs, DHC4-DHC6, or DHC8-DHC12. However, the iTRAQ ratios of three inner arm DHCs, DHC2, DHC3, and DHC7, were consistently reduced in *ida8* (Table 1). To confirm the DHC defects by label free quantification, we excised gel bands containing the DHCs (400-500 kD) from several axoneme samples, digested them with trypsin, and analyzed them by MS/MS and spectral counting (Zhu et al., 2010; Wirschell et al., 2013; Bower et al., 2013; 2018). As shown in Figure 5, DHC2, DHC3, and DHC7 were reduced in *ida8-1* and restored to wild-type levels in the *FAP57-HA* rescued axonemes, consistent with the ratios observed by iTRAQ labeling. Previous studies have shown that DHC2 elutes in peak *d* and DHC7 in peak *g*, that DHC3 is a minor dynein located in the proximal portion of the axoneme and closely related to DHC7, and that both dynein *d* and *g* are located at the distal end of the 96 nm repeat (Yagi et al., 2009; Bui et al., 2012; Kollmar, 2016). Collectively, these observations strongly suggested that FAP57 targets or stabilizes the attachment of these dyneins to their unique binding sites in the 96 nm repeat.

**Figure 5.**
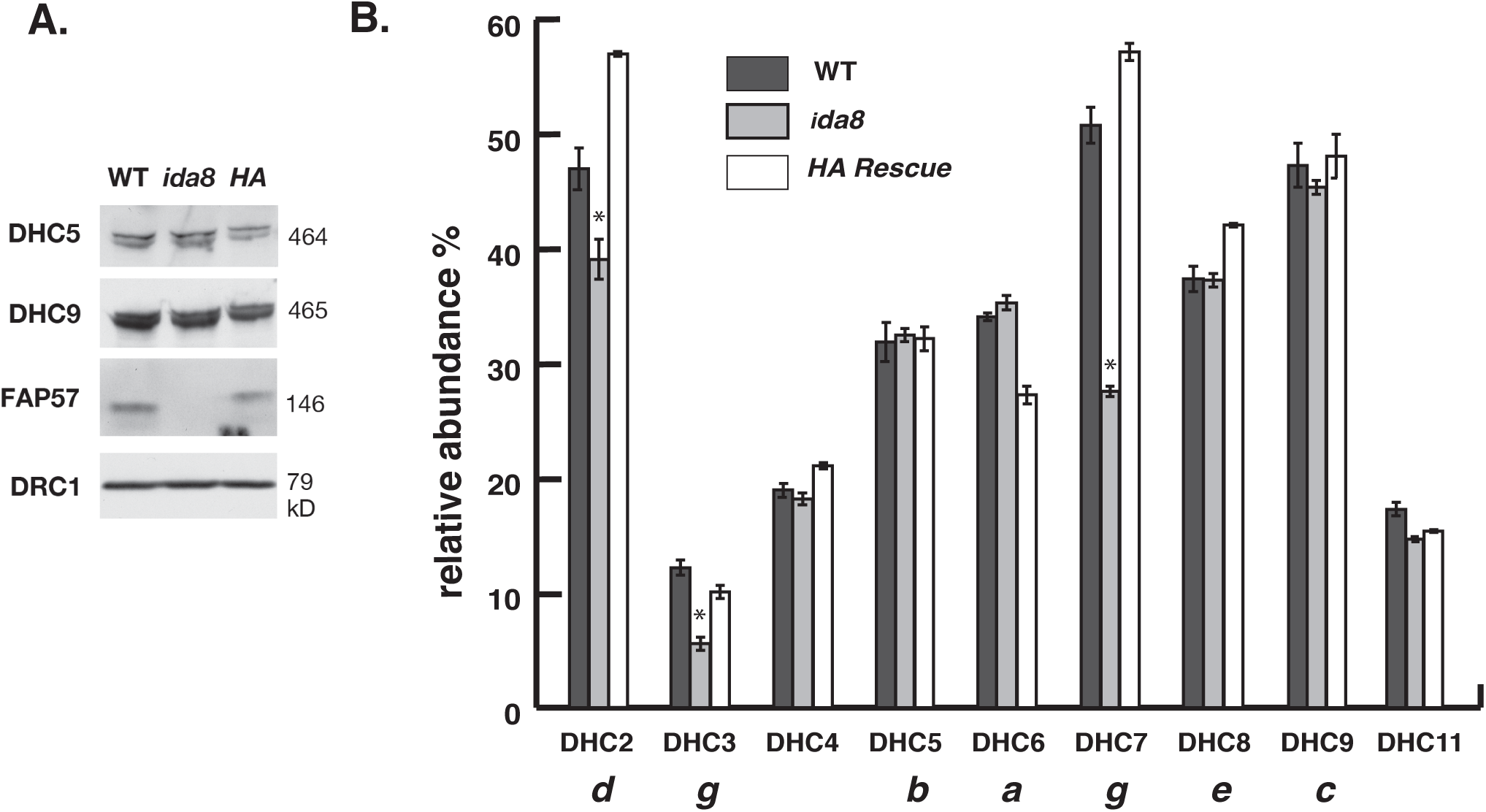
Mass spectrometry reveals defects in the assembly of a subset of inner arm DHCs in *ida8*. **(A)** A Western blot of axonemes from wild-type (WT), *ida8-1*, and an *ida8-1; FAP57-HA* rescued strain (*HA*) was probed with antibodies against two inner arm DHCs (DHC5, DHC9), FAP57, and DRC1. **(B)** The same samples were fractionated by SDS-PAGE, and the DHC region was excised and analyzed in triplicate by tandem MS/MS and spectral counting. The total counts for each DHC were expressed as a percentage of the total counts for the two I1 dynein DHCs.

### Cryo-electron tomography (cryo-ET) of ida8 reveals the complexity of defects in IDA structures

Given the complexity of the biochemical and structural defects observed in *ida8*, we analyzed wild-type, *ida8,* and *FAP57-*rescued axonemes by cryo-ET and subtomogram averaging to better resolve the defects in the structure of the IDAs. We also rescued *ida8* by transformation with N- and C-terminally SNAP-tagged versions of the *FAP57* gene to localize FAP57 more precisely and gain insight into the role of FAP57 in the targeting of IDAs. As shown in Supplemental Figure 6A and B, FAP57 polypeptides with either an N-terminal or a C-terminal SNAP tag were assembled into axonemes and restored the forward swimming velocity of *ida8* to near wild-type levels. Consistent with images obtained by TEM and 2D averaging (Figure 2C and King et al., 1994) but with higher resolution, the average of all *ida8* tomograms showed decreases in densities located in two distinct regions of the 96 nm repeat, one located at the distal end of the I1 dynein IC/LC complex (termed the I1-distal structure) and a second corresponding to the locations of the two distal dyneins, IDAs *g* and *d* (compare Supplemental Figure 6C, D with E, F). In addition, the density of IDA *b* appeared to be weaker in *ida8* relative to wild-type (Supplemental Figure 6C-F). The reduced densities were restored to wild-type levels in tomograms from the *FAP57* rescued strains (Supplemental Figure 6G-J).

**Figure 6.**
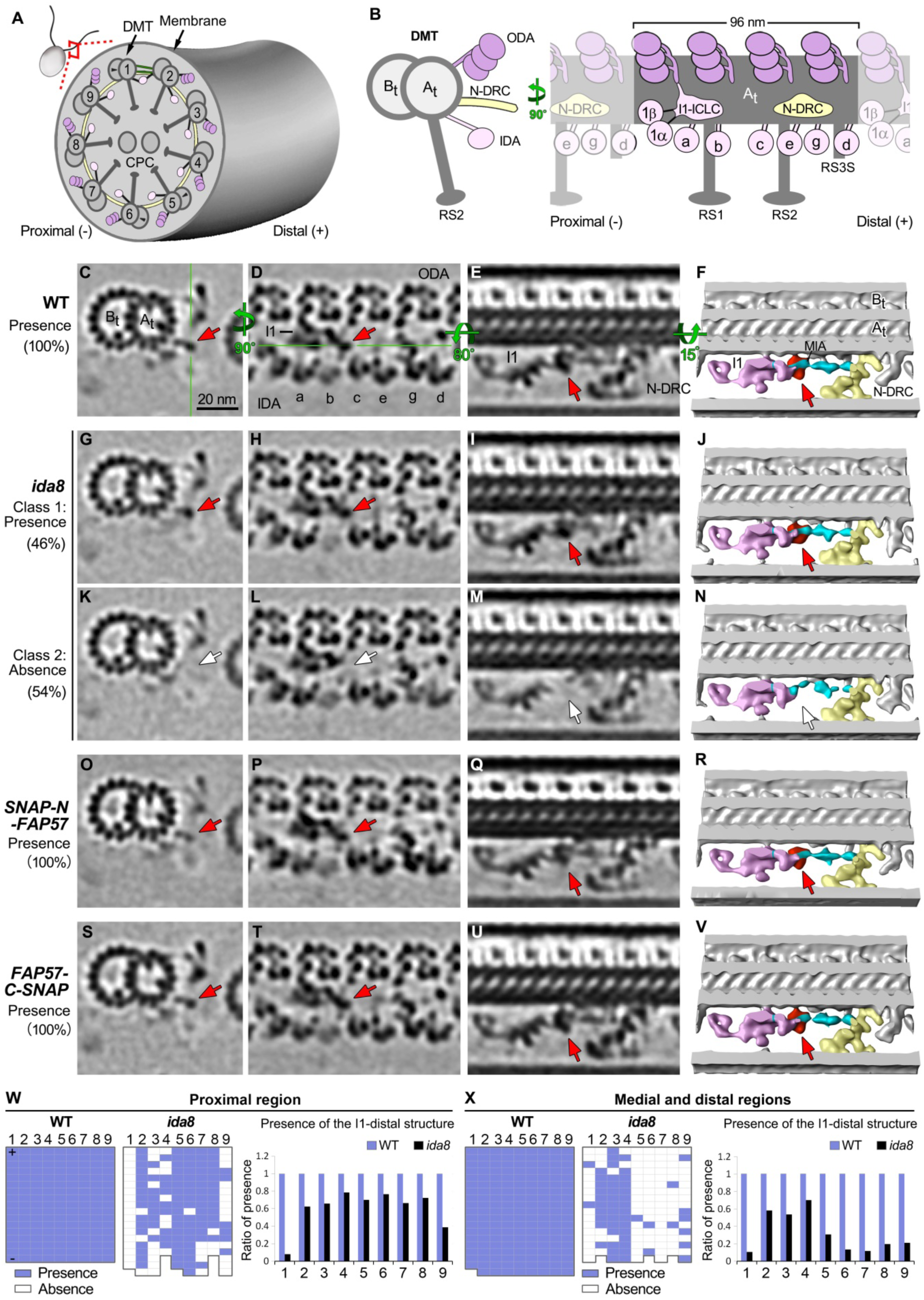
Cryo ET and class averaging of *ida8* reveal defects in a density located between I1 dynein and the MIA complex. **(A)** Diagram of a *Chlamydomonas* cell and one of its two flagella shown in cross-section with nine outer doublet microtubules (DMT1-9) surrounding the central pair complex (CPC). **(B)** Diagram of a single DMT shown in cross-section view (left) and longitudinal view from the perspective of the neighboring DMT (right), with the A- and B-tubules (A_t_ and B_t_). The DMT is built up by a series of 96 nm axonemal repeats, each of which contains four three-headed ODAs (dark pink) on top, three radial spokes or stumps (RS1, RS2, RS3S) at the bottom, and seven distinct IDAs (light pink) and the N-DRC (yellow) in the middle region. The two-headed I1 dynein is located at the proximal end, with its 1α and 1β dynein heads connected to the I1 IC/LC domain. The six single-headed IDAs (*a, b, c, e, g, d*) are attached to specific sites along the repeat. **(C-E)** Tomographic slices of the class average of the 96 nm repeat of WT axonemes showing the I1-distal structure in three different views: a cross section of 96 nm repeat through the I1-distal structure (C), a longitudinal section of the 96 nm repeat through the dyneins (D), and another longitudinal section that rotates 80 degrees (E). The green lines indicate the locations of the slices shown in next panel. Classification analysis showed that all WT repeats have the I1-distal structure. **(F)** Iso-surface renderings corresponding to image in (E), but with a small rotation for a better 3D view of the I1-distal structure (red) and its neighboring complexes: I1 dynein complex (pink), N-DRC (yellow), MIA complex (cyan). **(G-N)** Images of the two class averages of the 96 nm repeats in *ida8*. In Class 1 repeats (∼46%), the I1-distal structure was present (G-J, red arrows); in Class 2 repeats (∼54%), the I1-distal structure was missing (white arrows). **(O-V)** Images of *SNAP-N-FAP57* (O-R) and *FAP57-C-SNAP* (S-V) rescued axonemes, showing re-assembly of the I1-distal structure (red arrows). **(W and X)** The presence (blue grids) or absence (white grids) of the I1-distal structure in each 96 nm repeat was scored for individual DMTs in the tomograms taken from the proximal (W) or medial/distal regions (X) of the axoneme. Flagellar polarity is indicated by “+” and “–” ends. The WT dataset contained 5 proximal and 20 medial /distal tomograms, whereas the *ida8* dataset contained 16 proximal and 17 medial/distal tomograms. The averaged histograms on the right depict the ratio of repeats with the I1-distal structure relative to all repeats on the individual DMTs. Scale bar in (C) is 20 nm.

To better characterize the defects in *ida8* axonemes, we performed a classification analysis on each structure of interest. The proximal and medial/distal regions of the axoneme and the identities of DMTs 1-9 were determined by the presence of DMT specific features (Bui et al., 2012; Lin et al., 2012). These analyses not only precisely identified the structural defects in *ida8*, but it also correlated the defects with a specific region or DMT (Figures 6, 7). As shown in Figure 6, the I1-distal structure was present in 100% of the wild-type repeats (Figure 6C-F), missing in 54% (class 2) of the *ida8* repeats (Figure 6G-N), and recovered to 100% in the repeats from both rescued strains (Figure 6O-V). The missing structure is located on the distal side of the I1-dynein IC/LC complex, close to the location of the MIA complex (Yamamoto et al., 2013). Based on the iTRAQ results that detected WT levels of MIA proteins in *ida8* (Table 1), and the fact that some densities remained in this region in the *ida8* tomograms (Figure 6G-N), the I1-distal structure appears to be distinct from the densities associated with the MIA complex. Although defects in the I1-distal structure were observed on all DMTs of *ida8*, the classification analysis revealed that these defects were asymmetrically distributed both along the length of the axoneme and among the DMTs. More specifically, the I1-distal structure was most significantly reduced on DMTs 1 and 9 in the proximal region and on DMT1 and DMTs 5-9 in the medial/distal region (Figure 6W, X).

**Figure 7.**
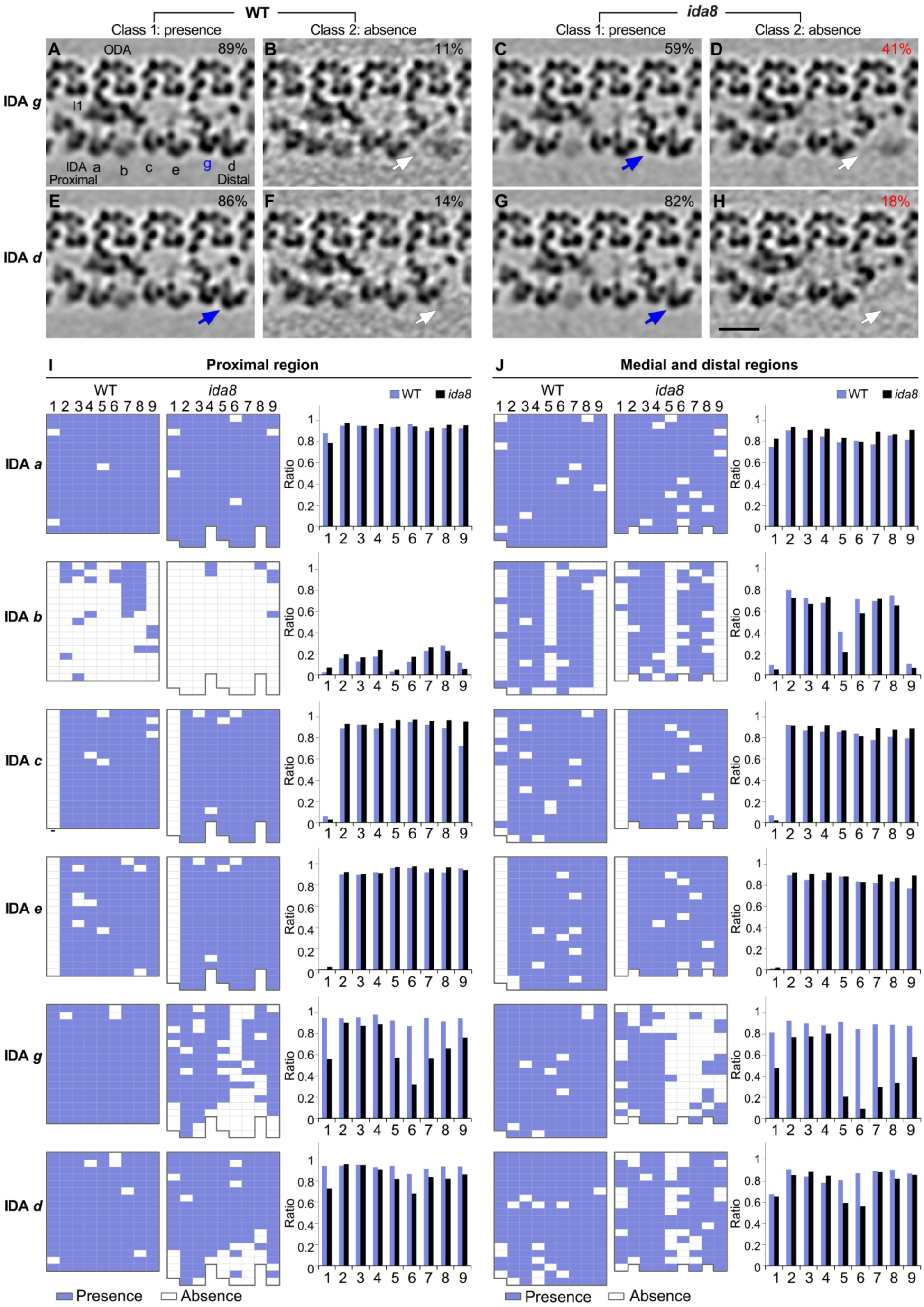
Cryo ET and class averaging reveal defects in the assembly of IDAs *d* and *g* on specific DMTs in *ida8.* The WT and *ida8* repeats were analyzed for the presence (Class 1) or absence (Class 2) of each single-headed IDA (*a, b, c, e, g, d*). **(A-H)** Tomographic slices of the class averages of the 96 nm repeat, with four ODAs on top and the single-headed IDAs at the bottom, showing the presence (Class 1, blue arrows) or absence (Class 2, white arrows) of the IDAs *g* and *d*, which are reduced in many *ida8* repeats. The percentage of subtomograms included in each class average is indicated. (See Supplemental Figure 7 for the class averages of all the single-headed IDAs.) **(I, J)** The presence (blue grids) or absence (white grids) of the indicated IDA in a 96 nm repeat was scored for each DMT (1-9) in tomograms taken from the proximal (I) or medial and distal (J) regions of the axoneme. The WT dataset contained 5 proximal and 20 medial /distal tomograms, whereas the *ida8* dataset contained 16 proximal and 17 medial/distal tomograms. The averaged histograms on the right depict the ratio of repeats with the indicated IDA relative to all of the repeats for each DMT (1-9). Classification analysis showed that the assembly of dyneins *a*, *b, c*, and *e* in *ida8* was not significantly different from wild-type. However, more *ida8* repeats lacked IDAs *g* and *d* than WT (D, H), and the defect in assembly was biased toward DMTs 1 and 5-9. Scale bar in (H) is 20 nm.

Classification analyses of the individual IDAs (*a, b, c, e, g,* and *d*) in the sub-tomograms revealed significant differences in the assembly of IDAs *g* and *d* in *ida8* axonemes, but no significant changes in the assembly of IDAs *a, b, c,* and *e* (Figure 7; Supplemental Figure 7). In addition, the defects in assembly of IDAs *g* and *d* were distributed asymmetrically, similar to defects in the I1-distal structure (Figures 6, 7). The defect in IDA *g* was more remarkable: 41% of *ida8* repeats lacked IDA *g* compared to 11% of wild-type repeats (Figure 7A-D). This difference was largely due to decreased assembly of dynein *g* on DMTs 5-8 and to a lesser extent on DMT1 and DMT9 (Figure 7I, J). IDA *d* was missing in 18% of the *ida8* repeats compared to 14% of wild-type repeats (Figure 7E-H). This small change was due to decreased assembly of dynein *d* on DMTs 1, 5, and 6 (Figure 7I, J). The missing IDAs *g* and *d* were restored to wild-type levels in the SNAP-tagged, *FAP57* rescued axonemes (Supplemental Figure 8).

The classification analyses initially also suggested a decrease in the assembly of IDA *b,* because IDA *b* was missing in 67% of the *ida8* repeats versus 52% of the wild-type repeats (Supplemental Figure 7E-H). However, the *ida8* dataset contained a higher proportion of tomograms from the proximal region of the axoneme (48%) than the WT dataset (20%). As reported previously, IDA *b* is only rarely seen in the proximal region of wild-type axonemes, and it is mostly absent from DMTs 1, 5, and 9 in the medial/distal region (Figure 7I, J, see also Bui et al., 2012, Lin et al., 2012). Therefore, after correlating the IDA *b* defective repeats with specific regions of the axoneme, no significant difference in the assembly of IDA *b* was found between *ida8* and WT (Figure 7I, J).

To verify the asymmetric distribution of the structural defects across the nine DMTs identified by classification analysis, we also performed DMT specific averaging on wild-type, *ida8*, and *FAP57* rescued axonemes. As shown in Supplemental Figure 8, the images of the DMT specific averages are consistent with the classification analyses. The densities of the I1-distal structure, and IDAs *g* and *d* on DMTs 1, 5-9 were clearly weaker in *ida8* than in WT, but they were not obviously different from WT on DMTs 2-4. In addition, the densities corresponding to the missing structures were all restored to wild-type levels in the SNAP-tagged, *FAP57* rescued axonemes (Supplemental Figure 8).

### Localization of FAP57 in the 96 nm repeat by SNAP-tagging and biotin-streptavidin-nanogold labeling

Because FAP57 has two distinct polypeptide domains, an N-terminal region with seven WD repeats and a C-terminal region with several coiled coil domains, we reasoned that these two domains might be arranged along the length of the DMT and facilitate the targeting or stabilization of the different structures missing in *ida8*. To test this hypothesis, we treated axonemes from the SNAP-tagged rescued strains with biotin and streptavidin-nanogold (+Au) and analyzed the tomograms for presence of additional densities that might reveal the locations of the N- and C-termini of the FAP57 polypeptide (Figures 8, 9). As shown in Figure 8A and B, averages of all repeats revealed an additional density in the *SNAP-N-FAP57* +Au sample that was located close to the site of the I1-distal structure previously identified as missing in *ida8* (Figure 8B, yellow arrows). To enhance the signal-to-noise ratio for detecting the streptavidin-nanogold label, we generated DMT specific averages using sub-tomograms from DMTs 6-8 (Figure 8C-J) and DMTs 2-4 (Figure 8K-R). The DMTs 6-8 averages clearly showed the missing I1-distal structure in *ida8* (Figure 8D, H, pink arrows), its recovery in the *SNAP-N-FAP57* strain (Figure 8E, F, red arrows), and the presence of an additional density in the streptavidin-gold-treated sample (Figure 8F, J, yellow arrows). Consistent with the classification results (Figure 7; Supplemental Figure 7), the DMTs 2-4 averages showed no obvious structural defect in *ida8* and no additional density in the streptavidin-gold-treated sample (Figure 8K-R). These results confirm the asymmetric distribution of the structural defects across the nine DMTs of *ida8* and suggest that the N-terminal portion of FAP57, which contains multiple WD repeats, contributes to the assembly of the I1-distal structure.

**Figure 8.**
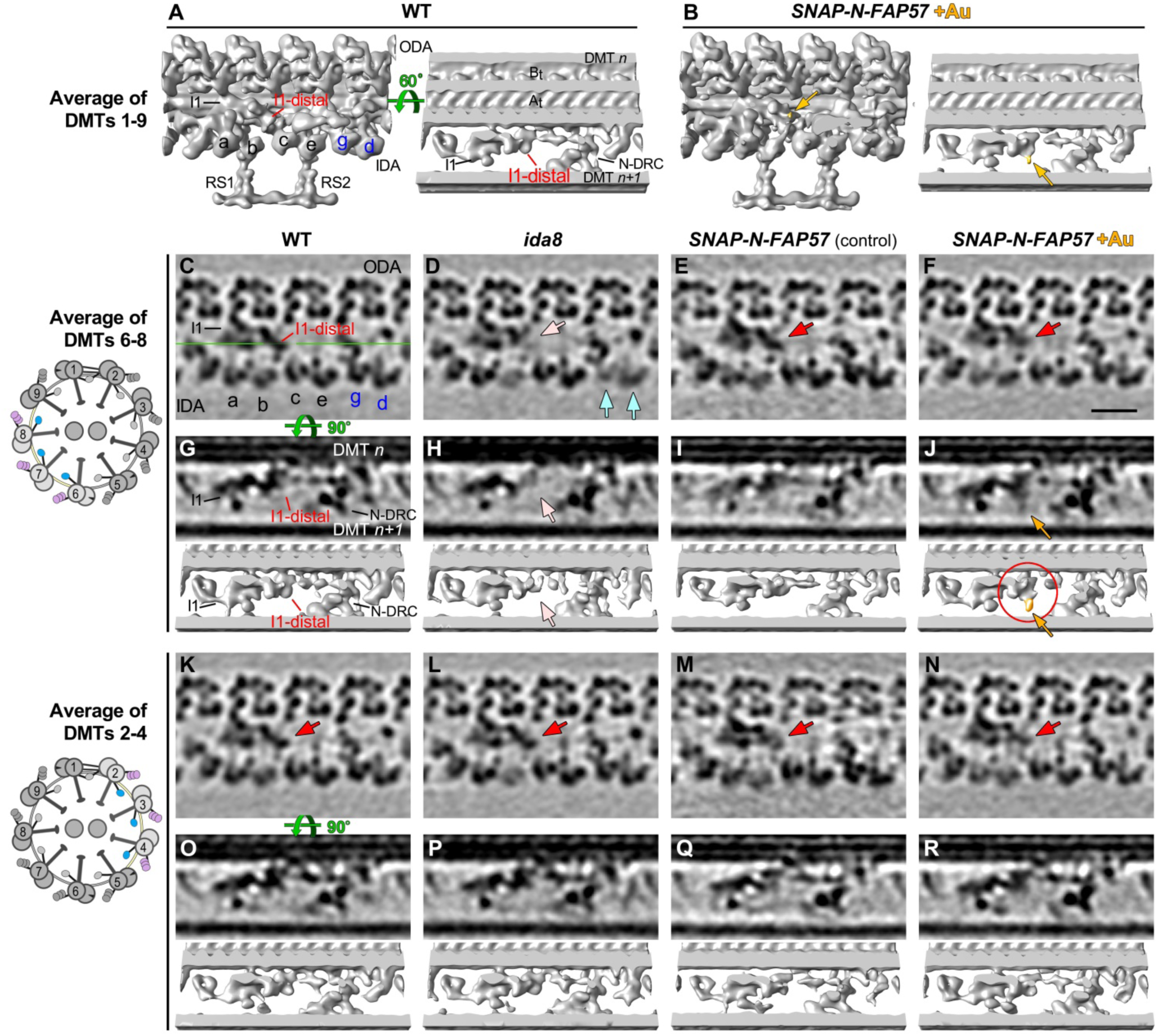
Streptavidin-gold labeling, cryo ET, and DMT specific averaging reveal the location of the N-terminus of FAP57. **(A, B)** Iso-surface renderings of the averaged 96 nm repeats from wild-type axonemes (A) or from streptavidin-gold labeled axonemes from *SNAP-N-FAP57* rescued strain (B). A new density was detected on the I1-distal structure in the gold-labeled rescued axonemes, as highlighted by yellow arrows. (**C-F**) Tomographic slices of the averaged 96 nm repeats of DMTs 6-8 from WT (C), *ida8* (D), *SNAP-N-FAP57* control (E), or gold-labeled *SNAP-N-FAP57* axonemes (F; +Au). The *SNAP-N-FAP57* control sample is different from the gold-labeled *SNAP-N-FAP57* sample because the BG-(PEG)12-Biotin was omitted in the control during the labeling procedure. A diagram of an axoneme cross section is shown on the left, with DMTs 6-8 highlighted in color. Defects in the I1-distal structure (pink arrows) and IDAs *g* and *d* (light blue arrows) were clearly visible as weaker densities in *ida8* (D). These densities were restored in both of the rescued samples (red arrows; E and F). (**G-J**) Tomographic slices (top) and isosurface renderings (bottom) of averaged 96 nm repeat of DMTs 6-8 from WT (G), *ida8* (H), *SNAP-N-FAP57* control (I), or gold-labeled *SNAP-N-FAP57* axonemes (J). The location of the tomographic slice is indicated by a green line in (C). The red circle in (J) highlights the new density on I1-distal structure observed in the gold-labeled sample. This density is noticeably larger than that seen in the average of all nine DMTs in (B). (**K-R**) Images of averaged 96 nm repeats of DMTs 2-4 from WT (K, O), *ida8* (L, P), *SNAP-N-FAP57* control (M, Q), or gold-labeled *SNAP-N-FAP57* axonemes (N, R). The I1-distal defect was hardly visible in averages from DMTs 2-4 of *ida8* (L, P), and a new density was not clearly visible in the gold-labeled axonemes (N, R). Scale bar in (F) is 20 nm.

**Figure 9.**
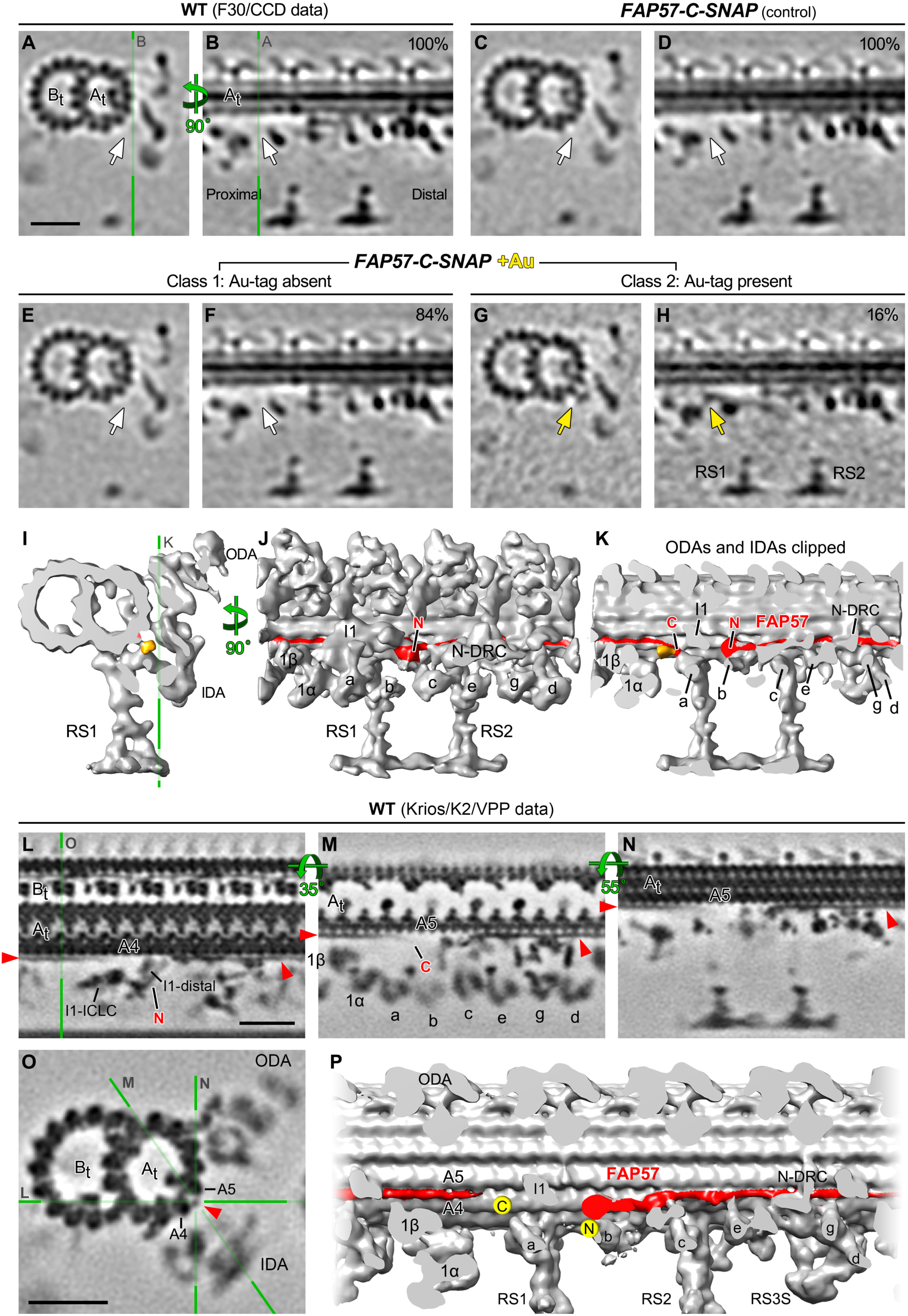
Streptavidin-gold labeling, cryo ET, and class averaging reveal the location of the C-terminus of FAP57, and higher resolution average directly visualizes the candidate density of FAP57. (A-H) Tomographic slices of class averages of the 96 nm repeat from WT axonemes (A, B), *FAP57-C-SNAP* control (C, D), or gold-labeled *FAP57-C-SNAP* axonemes (E-H; +Au). BG-(PEG)12-Biotin was omitted during the labeling protocol in the FAP57-C-SNAP control sample in (C, D). The averaged repeats are shown in cross-section (left) and in longitudinal section at a plane close to the surface of the A-tubule (right). The locations of tomographic slices are indicated by green lines. The percentage of sub-tomograms included in each class average is indicated. A new class (Class 2) was identified in the gold-labeled *FAP57-C-SNAP* axonemes in (G, H). Class 2 contained a novel density (gold arrows) located close to the surface of the A-tubule and proximal to radial spoke 1 (RS1) and the IC/LC complex of the I1 dynein. Such novel density was not observed in the control (C, D) or Class 1 averages (E, F, white arrows). (I-K) Isosurface rendering of the 96 nm repeat from the Class 2 axonemes viewed in cross-sectional (I) and longitudinal (J, K) orientations. The ODAs and IDAs were clipped in (K) for better visualization of the novel density. The clipping plane is indicated by the green line in (K). The predicted arrangement of the FAP57 polypeptide on the surface of A-tubule is shown in red. The proposed locations of the N- and C-termini of FAP57 are denoted based on the streptavidin-gold labeling of *SNAP-N-FAP57* (Figure 8) and *FAP57-C-SNAP* axonemes (this figure), respectively. (L-O) Longitudinal (L-N) and cross-sectional (O) tomographic slices of the 96 nm repeat from WT axonemes that have significantly improved resolution (1.8 nm, 0.5 criteria of FSC). The green lines with letters indicate the locations of the slices shown in corresponding panels. At this significantly improved resolution, a filamentous structure that extends from the I1-distal structure (L), attaches on protofilaments A4 and/or A5 of the A-tubule, and runs along the A-tubule towards the distal side was clearly visible (L-O, red arrowheads). The location of this structure coincides with the location of FAP57 predicted in (J, K), suggesting this structure as a compelling candidate of FAP57. (P) Iso-surface rendering of the higher resolution average in longitudinal orientation. The candidate density of FAP57 is highlighted in red and the proposed locations of its C- and N-termini were denoted by two yellow dots. Scale bars in (A), (L) and (O) are 20 nm; (A) valid for (A-H), (L) valid for (L-N).

The tomograms obtained from *FAP57-C-SNAP* axonemes showed recovery of structures missing in *ida8*, similar to the *SNAP-N-FAP57* axonemes (Figure 6 O-V, Supplemental Figures 6I-J, 8), but an additional density associated with the streptavidin-nanogold labeling of the C-terminal SNAP tag was much harder to detect than for the N-SNAP tag. These results may indicate that the C-terminus of FAP57 is buried within a complex of other proteins and less accessible to bind the streptavidin-gold in intact axonemes. To increase the signal-to-noise ratio for detection of the C-terminally labeled region of FAP57, we turned again to classification of the sub-tomogram averages (Figure 9A-K). This approach identified an additional density in a small subset (∼16%) of sub-tomograms (Figure 9E, F, I-K). This additional density was located close to the surface of the DMT, proximal to the base of RS1, near the base of the I1/*f* dynein and dynein *a*.

### Direct visualization of the candidate density of FAP57 with recent hardware and software advances for cryo-ET

Aiming at directly resolving the FAP57 structure in 3D, we applied recent hardware and software advances for cryo-ET and reanalyzed the 96 nm repeat in WT axonemes (Materials and Methods). We further improved the resolution of the averaged 96 nm repeat from 3-4 nm to 1.8 nm (0.5 criteria of FSC, Supplemental Table 5). The significantly higher resolution allowed clear visualization of an extended filamentous structure as a candidate for the FAP57 protein (Figure 9L-P). This filamentous structure extends from the I1-distal density, near the site of the N-terminus of FAP57, to the outer cleft of protofilaments A4-5 (Figure 9L), from where it runs parallel to the A-tubule towards the distal end of the 96 nm repeat, making attachments to multiple structures such as the N-DRC and tail domains of IDA *g* and *d* (Figure 9L-P). The filament then extends into the next 96 nm repeat and becomes slightly curved, with a weaker density at the end near the base of IDA *a,* close to the site where the C-terminus of FAP57 was localized (Figure 9M). This weaker density may be a reflection of the greater positional flexibility of the C-terminal region of FAP57, which is predicted to be a low complexity region that is potentially disordered. The flexibility of the FAP57 C-terminus in combination with the crowded molecular environment around the base of both the I1/*f* dynein and IDA *a* might explain the lower labeling efficiency with streptavidin-gold (Figure 9E, F).

## Discussion

### The BOP2/IDA8 locus encodes a conserved polypeptide required for stabilizing the binding of a subset of IDAs

Our study of the *bop2/ida8* mutations has identified a new sub-complex that contributes to the organization of IDAs within the 96 nm repeat. The three *ida8* alleles and *bop2-1* are null mutations that fail to assemble FAP57 into the axoneme (Figures 1, 2). Transformation with WT or epitope-tagged versions of FAP57 rescues the motility defects and restores the missing proteins (Figures 2, 3, 5). FAP57 is a highly conserved polypeptide containing an N-terminal region with multiple WD repeats and a C-terminal region with multiple coiled coil domains (Figure 2). Interestingly, closely related orthologues known as WDR65/CFAP57 have been identified in other organisms with motile cilia, but only in those species that assemble IDAs (Table 1; see also Nevers et al., 2017). The loss of FAP57 in *Chlamydomonas* is correlated with the absence of a second highly conserved WD repeat protein, FAP337, and reduced assembly of inner arm DHC2, DHC3, and DHC7 (Tables 1, 2, Figure 5; Supplemental Figure 5). The polypeptide defects observed in *ida8* are distinct from those described in other motility mutants (Supplemental Figure 3). Taken together, the results suggest that FAP57 and FAP337 form a distinct sub-complex that is required to stabilize the binding of specific IDAs at the distal end of the 96 nm repeat.

Quantitative mass spectrometry using both iTRAQ labeling and label-free spectral counting has shown that two other proteins, FAP331 and FBB7, are elevated in *ida8* but restored to WT levels in a rescued strain (Table 1, 2). Both proteins share a similar overall structural organization of N-terminal, WD repeat domains and C-terminal, coiled coil domains with FAP57 (Supplemental Figure 5). In particular, FBB7 shares significant sequence homology with FAP57 (Table 1). These observations are reminiscent of earlier studies on the *ida5* mutants, in which mutations in the conventional *Chlamydomonas* actin gene (*IDA5*) were offset by increased expression of a novel actin-related protein (*NAP1*) (Kato-Minoura et al., 1997, 1998). Actin is an IC subunit of all single-headed IDAs, and *ida5* mutations result in the failure to assemble IDAs *a, c, d,* and *e*. However, NAP1 substitutes for the missing actin subunit in IDAs *b* and *g* and permits their assembly in *ida5* mutants. Similar changes in expression were also observed with the redundant I1 tether head subunits FAP43 and FAP244 in *fap43* mutants (Fu et al., 2018). One possibility is that FBB7 may partially compensate for the absence of FAP57 and stabilize the binding of the remaining IDAs in the *ida8* mutant. Identification of a *fbb7* mutant will be required to test this hypothesis.

### FAP57 forms an extended structure within the 96 nm repeat that interacts with multiple regulators of IDA activity

Several lines of genetic, biochemical, and structural evidence suggest that FAP57 extends nearly the full length of the 96 nm repeat and interacts with multiple regulators of IDA activity. The first *fap57* mutation, *bop2-1*, was originally isolated as extragenic suppressor of the paralyzed flagellar mutant *pf10* (Dutcher et al., 1988). Little is known about the *PF10* gene product or the identity and location of polypeptides that might be altered in *pf10* axonemes. However, the gene product of another *pf10* suppressor, *BOP5*, has been identified as the I1 subunit IC138 (Hendrickson et al., 2004; VanderWaal et al., 2011). IC138 is a WD repeat protein within the IC/LC complex at the base of I1 dynein (Bower et al., 2009; Heuser et al., 2012). Given that two *pf10* suppressors have been linked to WD repeat proteins associated with IDAs, *pf10* may be directly or indirectly associated with defects in the coordination or regulation of IDAs.

Biochemical studies have also identified potential interactions between FAP57- and I1 dynein-associated proteins. Yamamoto et al. (2012) described FAP57 as one of a few polypeptides that co-immunoprecipitated with the FAP100 subunit of the MIA complex, the IC138 and two DHCs of I1 dynein, and the FAP44 subunit of the I1 dynein tether head after chemical cross-linking. FAP57 has also been linked to the FAP44 subunit of the I1 tether head in *Tetrahymena,* based on proximity labeling (Urbanska et al., 2018). We found that FAP57 co-elutes with the I1 tether head subunits during FPLC fractionation (Figure 4, Supplemental Table 4). Although none of these proteins require FAP57 for assembly into the axoneme (Table 2) nor vice versa (Supplemental Figure 3; see also Urbanska et al., 2018), collectively the data suggest that FAP57 is located in close proximity to both I1 dynein and the MIA complex.

Comparison of WT and mutant axonemes by TEM and cryo-ET has provided even more compelling evidence for a direct physical interaction between FAP57, I1 dynein, and the MIA complex. Loss of FAP57 in *ida8* and *bop2* results in a defect in the assembly of a globular structure located just distal of the I1 dynein, at the site where the I1 IC/LC complex contacts the MIA complex (Figures 2D, 6). Rescue of *ida8* with a SNAP-tagged FAP57 followed by streptavidin-gold labeling confirms that the N-terminus of FAP57 is located within the I1-distal structure (Figure 8, 10). We propose that the WD repeat domains located within the first half of the FAP57 polypeptide interact with related domains in FAP337 to form at least part of the I1-distal structure.

**Figure 10.**
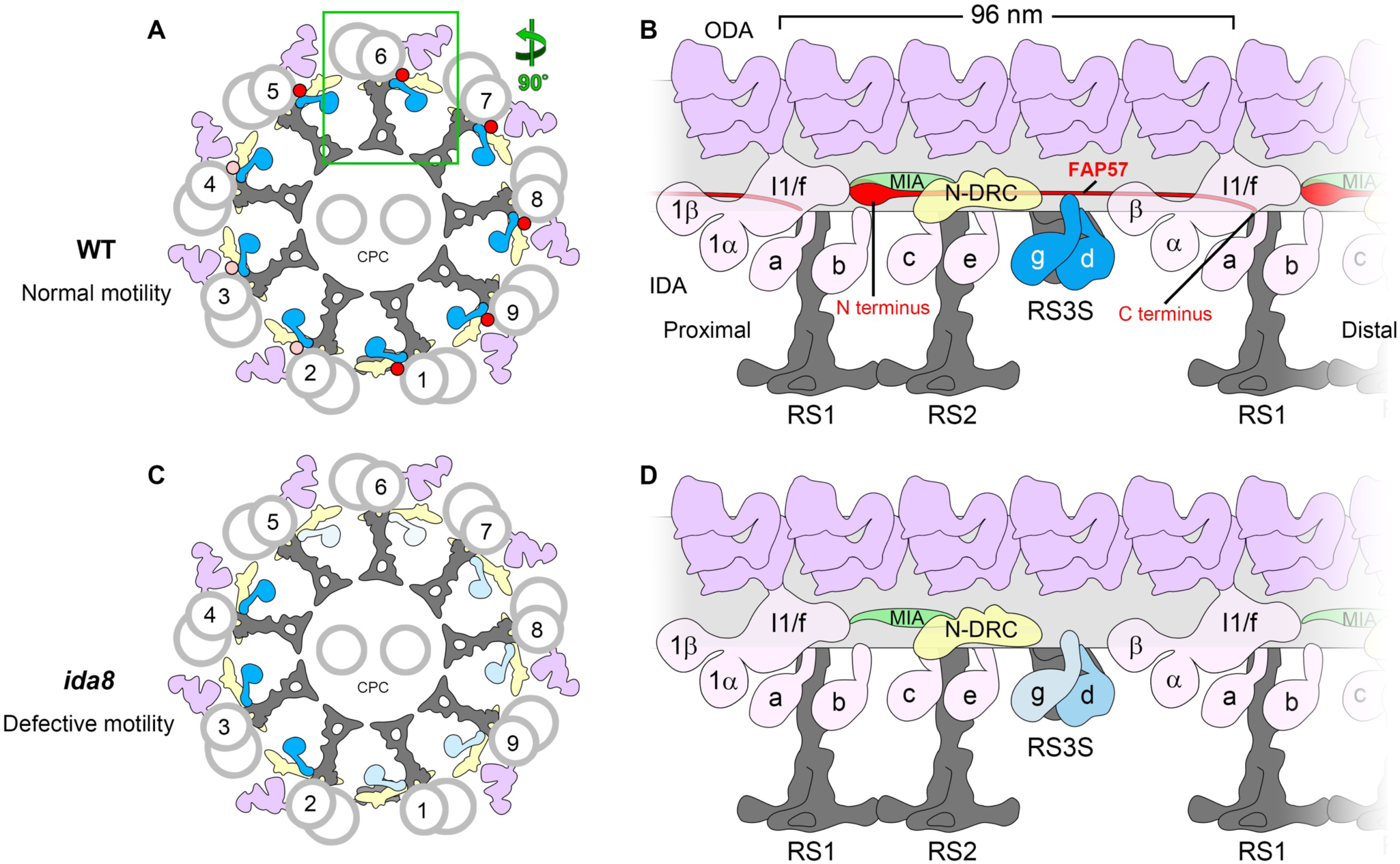
Model for the arrangement of FAP57 in the 96 nm repeat and its role in the assembly of certain IDAs. **(A)** Diagram of the cross-section of a WT axoneme showing the arrangement of the DMTs and the proposed asymmetric distribution of FAP57 across the nine DMTs: most FAP57 located on DMTs 1, 5-9 (red dots), while a small number of FAP57 are also located on DMTs 2-4 (pink dots). **(B)** Diagram of the longitudinal view of a WT DMT showing the proposed location of FAP57. The N-terminal portion of FAP57 containing the WD repeat domains is proposed to form the more globular structure that is located distal to the IC/LC complex of the I1 dynein. The second half of FAP57 containing the coiled coil domains is proposed to extend along the surface of the DMT, passing through or adjacent to the bases of IDAs *g* and *d*, and then extend further, with its C terminus located close to the base of RS1. The IDAs *g* and *d* are highlighted in blue. Note that FAP57 is proposed to contact multiple structures implicated in the regulation of IDAs, including the IC/LC complex of the I1 dynein, the MIA complex, the N-DRC, and IDAs *g* and *d*. **(C)** Diagram of the cross-section of an *ida8* axoneme showing the defects in the assembly of IDAs due to the loss of FAP57. In the absence of FAP57, fewer IDAs are assembled on DMTs 1, 5-9 (light blue) than on DMTs 2-4 (blue). **(D)** Diagram of the longitudinal view of an *ida8* DMT showing the proposed role of FAP57 in stabilizing the assembly of specific IDAs. In the absence of FAP57, the assembly of IDA *d* is only slightly reduced but IDA *g* is significantly reduced. The observed increase in FBB7 may compensate in part for the absence of FAP57. The levels of IDA *d* and *g* are shown by the intensity of the blue labels, with the lighter blue hues indicating less dynein present. Other labels: ODA, outer dynein arm; IDA, inner dynein arm; N-DRC, nexin-dynein regulatory complex; CPC, central pair complex; RS3S, radial spoke 3 stand-in.

Identifying the location of the C-terminal half of FAP57 (residues 623-1316) has been much more challenging. This region contains an extended coil-coil domain (residues 644-1188) followed by a more variable low complexity domain at the C-terminus (residues 1206-1314) (Figure 2). However, the defects in assembly of IDAs at the distal end of the 96 nm repeat observed in *ida8* (Figure 7, Supplemental Figures 6, 7, and 8) suggest that the C-terminal half of FAP57 extends from the I1-distal structure and runs close to the surface of the DMT, past the second RS (RS2) and N-DRC, to the sites of attachment for IDAs *g* and *d* (Figure 10). Moreover, rescue of *ida8* with a FAP57-C-SNAP construct followed by streptavidin-gold labeling and class averaging reveals the presence of an additional density compared to the wild type structure that is located near the base of the I1 dynein and IDA *a* (Figure 9). We propose that the coiled coil domain extends beyond the N-DRC into the next 96 nm repeat and that the additional density identifies the location of the C-terminus of FAP57 (Figure 10). This arrangement would be consistent with the predicted length of the coiled coil region (Surkont et al., 2015; Truebestein and Leonard, 2016) and the observation that FAP57 stabilizes the attachment of IDA *g* and *d*.

The challenges in directly visualizing defects in the assembly of a thin filamentous structure on the surface of the DMT are not without precedent. Indeed, localization of the axonemal ruler proteins FAP59/FAP172 relied on the presence of large tags to mark the positions of specific sites, but the proteins themselves could not be directly visualized by conventional cryo-ET (Oda et al., 2014). However, our recent efforts to improve resolution by combining cryo-ET with the methodologies used in single particle cryo-EM have yielded new images of axoneme structures with significantly greater structural detail (Song et al., BioRxiv). This approach made it possible to resolve the FAP59/FAP172 axonemal ruler as a filamentous structure located between protofilaments 2 and 3 of the A-tubule, at the base of the RSs in *Tetrahymena* axonemes (Song et al., BioRxiv). This study also identified another filamentous structure between protofilaments 4 and 5, at the base of several single-headed IDAs. This structure, named the inner arm ruler, is identical to the extended structure identified in our high-resolution average of the *Chlamydomonas* axonemal repeat (Figure 9L-P), with a location that coincides with the location of FAP57 predicted by our labeling results (Figures 9, 10).

### Asymmetry of IDA defects in both bop2 and ida8

A consistent feature of the *bop2/ida8* phenotypes is the asymmetry of IDA defects around the circumference of the axoneme, i.e. among the DMTs (King et al., 1994; Figures 7, 10; Supplemental Figure 8). The earlier study of *bop2-1* axonemes by TEM and 2D averaging indicated that defects in IDA assembly were limited to DMTs 5, 6, 8, and 9 (King et al., 1994). Here we used cryo-ET, doublet-specific and class-averaging focused on individual IDAs to show that loss of FAP57 in *ida8-1* impacts the assembly of IDA *g* on DMTs 5-8 and to a lesser extent on DMTs 1 and 9 (Figure 7). Given the nature of the mutations in both strains (Figures 1-5, Table 1), it is clear that although FAP57 is important for stabilizing the binding of IDA *g* (and to a lesser extent the binding of IDA *d*), it is not the only factor that specifies these dynein attachment sites. Indeed, we have previously noted defects in the assembly of IDA *g* and *d* in *n-drc* mutants (Heuser et al., 2009; Bower et al., 2013, 2018; Wirschell et al., 2013). As mentioned above, the FAP57 related protein, FBB7, is increased in *ida8* axonemes. FBB7 may also contribute to the targeting or stabilization of IDAs on specific DMTs.

Asymmetric defects in the assembly of dynein arms are not unique to *bop2/ida8* mutants. For example, a previous study of the *sup-pf2* mutations in the *γ* DHC revealed reduced assembly of ODAs on DMTs 3, 6-9 relative to DMTs 2, 4-5 (Rupp et al., 1996). Reduced assembly of ODAs on DMTs 6-9 was also seen in *oda4-s7* and *oda2-t*, two DHC truncation mutants (Liu et al., 2008). More recently, we observed reduced assembly of IDA *b* on DMTs 2-4 in the *pacrg* mutant; PACRG is one component of the inner junction between the A- and B-tubules of the DMT (Dymek et al., 2019). The factors that lead to the destabilization of dyneins on specific DMTs in these mutants are not well understood, but they may be related to the forces experienced by different DMTs during axonemal bending (Liu et al., 2008).

The asymmetry of dynein activity is believed to be essential for the generation of ciliary and flagellar motility. Indeed, a recent study of actively beating sea urchin sperm flagella revealed functionally distinct dynein conformations that alternated in a bend-direction specific manner between DMTs on opposite sides of the flagellum, i.e. between DMTs 2-4 and DMTs 7-9 (Lin and Nicastro, 2018). However, the mechanism(s) that switch the dynein activities during flagellar beating are unclear. The asymmetric distribution of FAP57 and its impact on the assembly of IDAs on specific DMTs suggests that FAP57 may be one component of the molecular mechanisms that regulate the pattern of dynein activity in beating cilia.

### Function of FAP57/WDR65 in ciliary motility in other species

Although several studies have identified FAP57/WDR65 as a conserved component of motile cilia and flagella (Broadhead et al., 2006; McClintock et al., 2008; Arnaiz et al., 2010; Sigg et al., 2017; Blackburn et al., 2017), functional studies on *fap57/wdr65* mutations are very limited. One report identified a missense mutation in WDR65 in a patient with Van der Woude syndrome, a cleft palate disorder, but this work did not establish a clear connection between the mutation and the cleft palate defects (Rorick et al., 2011). The *Drosophila* orthologue of FAP57/WDR65, *CG4329*, is expressed in the chordotonal neurons, which contain 9+0 cilia critical for the auditory response, and mutations in *CG4329* lead to moderate hearing impairment (Senthilan et al., 2012). The chordotonal neurons express ODAs, IDAs, and several dynein regulators, and dynein mutations also lead to hearing defects (Senthilan et al., 2012; Karak et al., 2015; zur Lage et al., 2019). Little is known about the motility of the 9+0 cilia in the chordotonal neurons, but the dyneins are thought to act as adaptation motors that amplify mechanical input (Senthilan et al., 2012; Karak et al., 2015). Whether *CG4329* regulates either the assembly or motility of dyneins in these 9+0 cilia remains to be determined. *CG4329* is also found in the *Drosophila* sperm proteome (Wasbrough et al., 2010), but a role in sperm motility has not yet been identified. However, the *Drosophila* orthologue of FAP337, *WDY* or *CG45799*, is expressed in the male reproductive tract, and maps to region of the Y chromosome that contains the male fertility factor *kl-1* (Vibranovski et al., 2008). Collectively these data hint at a potential role for WDR65 in axonemal motility in *Drosophila*. FAP57/WDR65 is also highly conserved throughout vertebrate species, and in humans, FAP57 is especially abundant in testes, respiratory tissue, and fallopian tubes (Fagerberg et al., 2014; Blackburn et al., 2017). We suggest that *FAP57/WDR65* should be screened as a candidate gene for those patients with PCD whose mutations have not been identified in the more commonly known PCD loci.

## Materials and Methods

### Culture conditions, genetic analyses, and strain construction

Strains used in this study (Supplemental Table 1) were maintained on Tris-acetate phosphate (TAP) medium, but occasionally resuspended in liquid minimal medium or 10 mM Hepes, pH 7.6, to facilitate flagellar assembly and mating. The three strains described here, *ida8-1* (59c2), *ida8-2* (45g11), and *ida8-3* (47d7), were isolated by transformation of a *nit1-305* strain with the plasmid pMN54 encoding the *NIT1* gene (Tam and Lefebvre, 1993; Mitchell and Sale, 1999). To verify that the motility defects were caused by *NIT1* insertion, strains were backcrossed to *nit1-305* strains with wild-type motility (either L5 or L8), and ∼120 tetrad progeny were scored for co-segregation of their motility phenotypes and their ability to grow on selective medium (R-NO_3_) in the absence of added nitrate. Some strains were crossed into an arginine-requiring background (either *arg7-2* or *arg7-8*) for transformation or complementation tests in stable diploids. Haploid *arg* strains were maintained by addition of 50 μg/ml arginine-HCL to TAP medium, and stable diploids (*arg 7-2/arg 7-8*) were selected after mating by plating on arginine-free media. Transformants were selected by co-transformation with pARG7.8 (Debuchy et al., 1989) and plating on arginine-free media or co-transformation with pSI103 (encoding the *aphVIII* gene) (Sizova et al., 2001) and plating on media containing 10 μg/ml paromomycin. Double mutant strains were recovered from progeny of non-parental ditype tetrads and confirmed by their motility phenotypes and Western blot analyses. The strains used for generating the higher resolution average of the 96 nm repeat in a pseudo WT strain (Figure 9) include *cw15* (CC-4533), and the CP-mutant strains *fap76-1*, *fap81*, *fap92, fap216,* and *fap76-1; fap81,* in which the DMT structure is undistinguishable from wild type (Fu et al., 2019). The latter are insertional mutants that were obtained from the *Chlamydomonas* Library Project (CLiP; https://www.chlamylibrary.org; Li et al., 2016; 2019).

### Characterization of plasmid insertion sites and recovery of the IDA8 gene

Purification of genomic, phage, and BAC DNA, restriction enzyme digests, agarose gels, isolation of RNA, preparation of cDNA, PCR and RT-PCR reactions, and Southern and Northern blots were performed as previously described (Perrone et al., 2000, 2003; Rupp et al., 2001; Rupp and Porter, 2003; Bower et al., 2013). Genomic DNA from each strain and several tetrad progeny was digested with a series of restriction enzymes and probed on Southern blots with ^32^P labeled pUC119 DNA to estimate the number of plasmid insertions (Supplemental Figure 1C). The plasmid DNA co-segregated with a slow swimming phenotype in fifteen random progeny from a cross between *ida8-1* and *nit1-305*. To identify the sites of plasmid insertion, genomic DNA was isolated from each strain, digested with restriction enzymes to release vector and flanking genomic DNA, treated with T4 ligase to re-circularize the plasmid, and transformed by electroporation into *E. coli* DH5*α*. The resulting plasmids were purified, digested, blotted, and probed with both pUC119 and *NIT1* DNA to identify restriction fragments that contained only flanking genomic DNA. An ∼700 bp *Sau3AI* band was recovered and subcloned as flanking clone 1 (FC1). Southern blots of genomic DNA probed with FC1 confirmed the presence of a RFLP in *ida8-1*.

FC1 was used to screen a genomic phage library (Schnell and Lefebvre, 1993) by colony hybridization and plaque purification. DNA isolated from positive clones was restriction mapped, subcloned, and used to rescreen the library to extend the chromosome walk in both directions. Subclones were tested on Southern blots to determine the extent of DNA re-arrangement or deletion caused by each insertion event (Supplemental Figure 1D). Subclones were also tested on Northern blots of total WT RNA isolated before and after deflagellation to identify the number and locations of the transcription units in the region. Six phage clones spanning ∼40 kb were tested for their ability to rescue the motility defects by co-transformation of *ida8-1; arg7-2* with the plasmid pArg7.8 and selection on medium lacking arginine. Over 500 *Arg+* transformants were screened per clone, but none of the clones rescued the motility defect. Because the complete transcription unit might not be contained within a single phage insert, a BAC library was also screened (https://www.chlamycollection.org/product/bac/). Five positive BAC clones (16p22, 34a14, 35d14, 6h9, and 7h7) were recovered and restriction mapped, and together they formed a contig of ∼117kb. Four strains with wild-type motility were recovered out of 400 *Arg+* positive colonies following co-transformation with BAC clone 6h9 (∼1% rescued). An ∼12.7 kb genomic clone containing the full-length *IDA8* transcription unit was subcloned using *Eco*RV and *Nde*1, ligated into *SmaI* digested pUC119 after partial fill-in, and transformed into *E.coli* DH10b cells by electroporation to generate the plasmid p59c2. Co-transformation of *ida8-1; arg7-2* with p59c2 rescued the motility defects (9/131 transformants or 6.9% rescue). Because the *IDA8* gene was cloned prior to completion of the *Chlamydomonas* genome project, the genomic DNA and predicted cDNA sequences of the *IDA8* transcription unit were determined by PCR and RT-PCR (Supplemental Table 2). Sequence files were analyzed using the Sequencher (Gene Codes, Ann Arbor, MI) and MacVector (Apex, NC) software packages.

### Mapping of the IDA8 gene and identification of bop2-1 as an ida8 allele

To place the *IDA8* locus on the genetic map, a genomic fragment was used to identify an *EcoRI/XhoI* restriction fragment length polymorphism (RFLP) between two strains, 137c and S1-D2 (Gross et al., 1988). The fragment was then hybridized to a series of mapping filters containing DNA isolated from the tetrad progeny of crosses between multiply marked *C. reinhardtii* strains and S1-D2. The segregation of the RFLP was analyzed relative to the segregation of more than 42 genetic and molecular markers (Porter et al., 1996; Kathir et al., 2003). The *IDA8* sequence was linked to the genetic marker *pyr1* (PD:NPD:TT = 7:0:7, ∼49 map units) and a molecular marker for *α*2 tubulin (PD:NPD:TT = 26:0:2, ∼3.6 map units) on the left arm of Linkage Group IV, close to the predicted location of the *bop2-1* mutation (Dutcher et al., 1988). This distance was consistent with the later sequence assembly of Chromosome 4, with *α*2 tubulin (Cre04.g216850) at nucleotides 144030-147635 and FAP57 (Cre04.g217914) at nucleotides 370547-382328. To determine if *bop2-1* might be an *IDA8* mutation, genomic DNA and RNA were isolated from *bop2-1* and analyzed by PCR, RT-PCR, and DNA sequencing (Bower et al., 2013; 2018). Co-transformation of *bop2-1; arg7-8* with p59c2 rescued the motility defects (3/60 transformants or ∼5% rescued).

### Characterization of the IDA8 gene product and generation of a specific antibody

The predicted amino acid sequence was compared to predicted sequences found in several versions of the *Chlamydomonas* genome project (https://phytozome.jgi.doe.gov/pz/portal.html). Predicted domains were identified using programs available at https://www.expasy.org/proteomics and https://iupred2a.elte.hu/. To identify a peptide that could be used to generate a specific antibody, regions of the amino acid sequence were analyzed for antigenicity using MacVector. Potentially immunogenic peptides were searched against all sequences available in the genome project to increase the likelihood that the chosen peptide would be unique. Peptides were also searched against predicted amino acid sequences in other species to identify regions of high sequence conservation. The peptide NLRGHNGKVRSVAWSPDDSKL (corresponding to amino acid residues 460-480) was synthesized, conjugated to KLH, and used to immunize two rabbits (Research Genetics, Huntsville, Alabama). Immune sera were tested by ELISA and affinity purified against the peptide.

### Epitope tagging of FAP57

For epitope-tagging of the C-terminus of FAP57, a 1.2 kb *SpeI-EcoRI* fragment spanning the predicted stop codon was isolated from p59c2 and subcloned into pBluescript. A *NdeI* site was created in the stop codon (TAA to TAT) using the Quick Change II XL Mutagenesis kit (Stratagene) and primers listed in Supplemental Table 2. A triple-HA epitope tag was amplified from the plasmid p3HA (gift of Carolyn Silflow, University of Minnesota, Saint Paul, MN), and a SNAP tag was amplified from a codon optimized SNAP plasmid (Song et al., 2015) using primers with *NdeI* sites (Supplemental Table 2). Both tags were inserted into the *NdeI* site of the 1.2 kb *SpeI/EcoRI* fragment and sequenced to verify orientation and reading frame. The tagged fragments were reinserted back into the original p59c2 plasmid to make pFAP57-HA and pFAP57-C-SNAP. To tag the N-terminus of FAP57, a 453 bp genomic fragment spanning the start codon was removed from the original 59c2 plasmid using two unique restriction sites, *AfiII* and *AvrII.* A 1032 bp version of this sequence was synthesized with the SNAP tag sequence inserted prior to the start codon (GeneWiz, South Plainfield, NJ) and then ligated back into the original p59c2 plasmid to make pSNAP-N-FAP57. All constructs were verified by sequencing in both directions and then linearized with *SspI* prior to co-transformation into *ida8-1* or *bop2-1*. Rescue of motility defects by co-transformation with the epitope-tagged FAP57 constructs was similar to that seen with the wild-type gene (∼3-6% rescued). The predicted amino acid sequences of the epitope-tagged FAP57 polypeptides are shown in Supplemental Figure 2.

### Phase contrast and fluorescence microscopy and measurements of swimming velocity and microtubule sliding

Motility phenotypes were assessed by phase contrast microscopy using a 20x or 40x objective on a Zeiss Axioskop microscope. Initial measurements of swimming velocities were made from video recordings using a C2400 Newvicon camera and Argus 10 video processor (Hammamatsu Photonic Systems, Bridgewater, NJ) calibrated with a stage micrometer (Myster, et al., 1997; 1999; Perrone et al., 1998, 2000). More recent assays used a Rolera-MGi EM-CCD camera (Q-imaging, Surrey, BC, Canada) and Metamorph software (Molecular Devices, San Jose, CA) (VanderWaal et al., 2011; Bower et al., 2013, 2018; Reck et al., 2016). To compare flagellar waveforms, a pco.1200HS camera with Camware software (Cooke Corporation, Londonderry, NH) was used to capture high-speed images (500 fps) of motile cells and create videos at 30 fps (Dymek et al., 2019). Videos and montages were created in Image J (Schneider et al., 2012).

Measurements of microtubule sliding velocities in protease-treated axonemes were performed as previously described (Okagaki and Kamiya, 1986; Dymek et al., 2019). A Zeiss AxioSkop 2 microscope equipped for dark-field optics with a Plan-Apochromat 403 oil immersion objective lens with iris and an ultra-dark field oil immersion condenser was used for imaging. Images were captured and analyzed using an ORCA-Flash 4.0 V2 (Hamamatsu) camera and Nikon NIS Elements Advanced Research Software (Tokyo, Japan). Data are presented as the mean +/- SEM. At least 3 independent experiments were performed for each strain. The Student’s *t*-test was used for comparisons between different strains.

Cells were fixed for immunofluorescence microscopy using ice-cold methanol (Sanders and Salisbury, 1995), stained with a rat monoclonal antibody to HA (Roche clone 3F10) and an Alexafluor-488 conjugated secondary antibody (Molecular Probes, Eugene, OR), and processed as previously described (Bower et al., 2013, 2018; Reck et al., 2016). Images were collected on a Zeiss Axioscop using a 100x/1.3 NA Plan Neuofluor objective, a CoolSnap ES CCD camera (Photometrics, Tuscon, AZ), and Metamorph software. Selected images were cropped, rotated, and labeled in Image J and Adobe Photoshop (San Jose, CA).

### Isolation and fractionation of axonemes, SDS-PAGE, and Western blot analyses

*Chlamydomonas* whole cell lysates, isolated flagella, and de-membranated axonemes were prepared as previously described (Witman, 1986; Bower et al., 2013, 2018; Reck et al., 2016) using 0.1-1.0% Nonidet-P-40 to remove membrane and matrix proteins. Purified axonemes were resuspended in HMEEN (10 mM Hepes, pH 7.4, 5 mM MgSO4, 1 mM EGTA, 0.1 mM EDTA, 30 mM NaCl) plus 1 mM DTT and 0.1 μg/ml protease inhibitors (leupeptin, aprotinin, pepstatin), and extracted with HMEEN containing 10mM MgATP, 0.6M NaCl, 0.2M NaI, 0.4M NaI, or 0.6M NaI. The 0.6M NaCl extracts (containing most of the axonemal dyneins and FAP57) were dialyzed against HMEEN, clarified by centrifugation, and fractionated on 5-20% sucrose density gradients (Bower et al., 2013; 2018) or diluted 10-fold and fractionated by Mono-Q ion-exchange FPLC chromatography (Gardner et al., 1994). Samples were separated on 5-15% polyacrylamide gradient gels and silver stained or transferred to Immobilon P and probed with different antibodies (Bower et al., 2013; 2018). Antibody sources and dilutions are listed in Supplemental Table 3.

### Preparation of samples for iTRAQ labeling and tandem mass spectrometry (MS/MS)

Isolated axonemes were washed in 10 mM Hepes pH 7.4 to remove salt, DTT, and protease inhibitors, then resuspended in 0.5 M triethylammonium bicarbonate pH 8.5 and processed for trypsin digestion and iTRAQ labeling as described in detail (Bower et al., 2013, 2018; Reck et al., 2016). Duplicate aliquots of axonemes (50-60 μg each) from each strain were reacted with 4-plex iTRAQ reagents (114-117, AB Sciex, Foster City, CA) to obtain two technical replicates per biological sample. The four labeled aliquots were mixed together and processed to remove excess trypsin, unreacted iTRAQ reagents, and buffer. The combined sample (containing two control aliquots with different iTRAQ labels and two mutant aliquots with different iTRAQ labels) was fractionated offline using high pH, C18 reversed phase chromatography (Reck et al., 2016). The column fractions were then further processed and loaded in 1-1.5 µg aliquots for capillary LC using a C18 column at low pH. The C18 column was mounted in a nanospray source directly in line with a Velos Orbitrap mass spectrometer (Thermo Fisher Scientific, Inc., Waltham, MA). Online capillary LC, MS/MS, database searching, and protein identification were performed as previously described (Lin-Moshier et al., 2013; Reck et al., 2016) using ProteinPilot software version 5.0 (AB Sciex, Foster City, CA) and the most recent version of the *Chlamydomonas* database (https://phytozome.jgi.doe.gov/pz/portal.html). The bias factors for all samples were normalized to *α* and *β* tubulin (Reck et al. 2016). The relative amount of protein in each aliquot was compared to that present in the control aliquot to obtain a protein ratio. The WT/WT or HA/HA ratios indicated the variability in labeling and protein loading between technical replicates of the same sample (typically less than 10% for all proteins). All iTRAQ experiments were repeated with a second set of samples for independent biological replicates. A total of 1060 proteins were identified at a 1% false discovery rate in the first experiment, and 1098 proteins were identified in the second experiment. The protein lists were filtered using a minimum of 6 peptides per protein, yielding 603 proteins from the first set of samples and 554 proteins from the second.

Because inner arm DHCs vary widely in abundance, purified axonemes from WT, *ida8-1*, and the *FAP57-HA* rescued strain were also fractionated by SDS-PAGE, stained briefly with Coomassie blue, and the DHC region was excised from the gel to improve the signal to noise for the DHCs (Bower et al., 2013). Following extraction and trypsin digestion, 3-5 replicates per sample were analyzed by MS/MS, and both the total number of peptides and total number of assigned spectra per DHC isoform were determined. The relative abundance of each DHC was estimated by spectral counting (Zhu et al., 2010) and expressed as a percentage of the total spectra identified for the 1-*α* and 1-*β* DHCs of the I1 dynein (Bower et al., 2013, 2018; Wirschell et al., 2013). Bands containing other proteins identified as significantly altered in the iTRAQ experiments were also analyzed by spectral counting to confirm the results with additional biological replicates. In addition, a subset of FPLC fractions was analyzed by MS/MS to identify polypeptides that co-fractionate with the FAP57 polypeptide. Samples were analyzed by SDS-PAGE, stained briefly with silver (Bower et al., 2013), and selected bands were excised from the gel and analyzed as described above (Supplemental Table 4).

### Thin section electron microscopy and image averaging

Isolated axonemes were fixed, resin-embedded, and sectioned for imaging by TEM as previously described (O’Toole et al., 1995; 2012). Cross-sectional and longitudinal views of axonemes were 2D averaged using the image processing software developed by the Boulder Laboratory for 3-Dimensional Microscopy and available at http://bio3d.colorado.edu/. To control for possible variations in staining, three to five biological replicates were processed for each sample. For averages of DMTs in cross-section, 183-659 DMTs were used to obtain a grand average for each strain. For averages of the 96 nm repeat in longitudinal section, individual averages containing multiple repeats were obtained for each axoneme, and 6-40 axonemes were analyzed for each strain to obtain a grand average (see Figure 2D).

### Cryo-electron tomography and image processing

The purified axoneme pellet was resuspended in HMEEK buffer (30 mM HEPES, pH7.4, 5 mM MgSO_4_, 1 mM EGTA, 0.1 mM EDTA, 25 mM KCl), and the suspension was directly used for cryo-sample preparation. For precise localization of the amino and carboxyl terminal ends of FAP57, streptavidin-nanogold labeling was performed on axonemes from strains rescued with SNAP-tagged versions of the FAP57 as previously described (Song et al., 2015). Briefly, 1 µl of 1 mM BG-(PEG)12-biotin (New England Biolabs; PEG linker available on request) was added to 200 µl of axonemes. A control sample was also prepared without added BG-(PEG)12-biotin. Both suspensions were incubated overnight at 4 °C, followed by three cycles of resuspension with HMEEK buffer and centrifugation at 10,000 g for 10 min at 4 °C. The axoneme pellets were resuspended in 200 µl of buffer, and then 5 µl of 1.4-nm-sized streptavidin-nanogold particles (strep-Au 1.4 nm, Nanoprobes, Inc) was added, and the two suspensions were incubated at 4 °C in the dark for 3 h with rotation. The samples were then diluted with 1 ml of HMEEK buffer, pelleted by centrifugation at 10,000 g for 10 min at 4 °C, carefully resuspended in 200 µl of HMEEK buffer, and used for cryo-sample preparation.

Cryo-sample preparation, cryo-ET and image processing were done as previously described (Nicastro et al., 2006; Nicastro, 2009; Heuser et al., 2009; Lin et al., 2014; Fu et al., 2018). Briefly, Quantifoil copper grids (Quantifoil Micro Tools, Jena, Germany) with a holey carbon film (R2/2, 200 mesh) were glow discharged for 30 seconds at −40 mA and loaded with 3 µl of axoneme sample and 1 µl of five-fold concentrated and BSA coated 10 nm colloidal gold (Sigma-Aldrich, St. Louis, MO) (Iancu et al., 2007). After brief mixing, grids were blotted from the back with Whatman No.1 filter paper for 1.5-3 seconds and plunge frozen in liquid ethane using a home-made plunger. Vitrified samples were cryo-transferred to a Tecnai F30 transmission electron microscope (ThermoFisher Scientific, Waltham, MA) for imaging. Single-axis tilt series of non-compressed, intact axonemes were acquired using the software package SerialEM (Mastronarde, 2005). Typically, 50 to 70 images were recorded at 13,500 fold magnification (∼1 nm pixel size) with −6 to −8 µm defocus while the specimen was tilted from about −65 to +65° in 1.5 to 2.5° increments. The microscope was operated in low dose mode at 300 keV and the cumulative electron dose for each tilt series was restricted to ∼100 e^-^/Å^2^ to minimize radiation damage. Electron micrographs were recorded digitally with a 2k x 2k CCD camera (Gatan) after passing a post-column energy filter (Gatan) in zero-loss mode with a slit width of 20 eV.

For higher resolution 3D structure of a pseudo wild-type axonemal repeat, vitrified axoneme samples from WT and CLiP mutants *fap76-1*, *fap81*, *fap92*, *fap216,* and a *fap76-1; fap81* double mutant were imaged by cryo-ET using a Titan Krios transmission electron microscope (Thermo Fisher Scientific, Hillsboro, OR) equipped with a K2 direct electron detection camera (Gatan, Pleasanton, CA) operated in counting mode, and a Volta-Phase-Plate (Danev et al., 2014). Tilt series were recorded at the magnification of 26,000 (∼5.5 Å pixel size) and −0.5 µm defocus using SerialEM with a dose-symmetric tilting scheme (Hagen et al., 2017). At each tilt angle, a movie stack of 15 frames was collected with a total exposure time of 6 seconds and an electron dose rate of 8 electrons/pixel/second. Motion correction of the frames was later performed on the movie stacks with a script extracted from IMOD (Kremer et al., 1996), and the resulting images of individual tilts were assembled into the tilt series.

3D tomograms were reconstructed from the recorded tilt series using fiducial alignment and weighted backprojection in IMOD. Sub-tomograms containing the highly repetitive 96 nm repeat units were further aligned and averaged with PEET (Nicastro et al., 2006), resulting in 3D structures with compensated missing wedge, reduced noise and thus increased resolution. For doublet-specific averaging, the nine DMTs were identified based on DMT-specific features (Bui et al., 2012; Lin et al., 2012), and repeats from individual DMTs were averaged. To further analyze structural defects that appeared heterogeneous or to identify the sites labeled with streptavidin-nanogold, classification analyses were performed on the aligned sub-tomograms using the PEET program (Heumann et al., 2011). Appropriate masks were applied to focus the classification analyses on specific regions of interest. Sub-tomograms containing the same structures were grouped into class averages. The structures were mapped onto their respective locations in the raw tomograms to determine the distribution of the different classes within the axonemes. The numbers of tomograms, sub-tomograms analyzed and the resolutions of the resulting averages are summarized in Supplemental Table 5. The resolution was estimated at the center of the DMT of axonemal repeat using the Fourier shell correlation method with a criterion of 0.5. The structures were visualized as 2D tomographic slices and 3D isosurface renderings using IMOD and UCSF Chimera (Pettersen *et al*., 2004), respectively.

### Online supplemental material

Fig. S1 describes the molecular characterization of the *ida*8 mutants, their motility phenotypes, and defects in axoneme structure observed by conventional transmission electron microscopy. Fig. S2 shows the predicted amino acid sequences of FAP57 and its epitope tagged variants. Fig. S3 depicts the distribution of FAP57 in axonemes from different *Chlamydomonas* motility mutants and in different flagellar extracts. Fig. S4 shows the motility phenotypes of *pf10, ida8, bop2* and the double mutants. Fig. S5 illustrates the predicted structural domains of the polypeptides that are altered in *ida8-1* axonemes. Fig. S6 demonstrates the rescue of *ida8-1* with the two SNAP-tagged *FAP57* constructs. Fig. S7 shows the longitudinal tomographic slices of the 96 nm repeat taken from class averages for the different IDAs in WT and *ida8-1.* Fig S8 shows the asymmetric distribution of structural defects in *ida8-1* axonemes and their recovery in SNAP-tagged FAP57 strains as revealed by DMT specific averaging.

Table S1 lists the strains used in this study. Table S2 lists the oligonucleotide primers used for RT-PCR and sequencing of *FAP57* transcripts in WT and *bop2-1*, and the oligonucleotide sequences used to generate epitope-tagged *FAP57* constructs. Table S3 lists the antibodies used in this study. Table S4 lists the proteins identified by MS/MS of the dynein peak *g* shown in Figure 4C. Table S5 summarizes the strains used for cryoET and corresponding image processing information. References specific to the supplemental material are also included.

Supplemental videos 1-8 show the waveforms of forward swimming cells as follows: Video S1 (WT), Video S2 (*ida8-1*), Video S3 (*bop2-1*), Video S4 (*ida8-1; FAP57-HA*), Video S5 (*bop2-1, FAP57-HA*), Video S6 (*pf10*), Video S7 (*ida8-1; pf10*) and Video S8 (*bop2-1; pf10*).

## Abbreviations

CSC: Calmodulin- and spoke-associated complex
CP: Central pair
CLiP: *Chlamydomonas* Library Project
DIC: Differential interference contrast
DHC: Dynein heavy chain
ET: Electron tomography
FAP: Flagellar associated polypeptide
HA: Hemagglutinin
IDA: Inner dynein arm
IC: Intermediate chain
IFT: Intraflagellar transport
iTRAQ: Isobaric tag for relative and absolute quantitation
LC: Light chain
N-DRC: Nexin-dynein regulatory complex
ODA: Outer dynein arm
*pf*: Paralyzed flagella
PEET: Particle Estimation for Electron Tomography
PCR: Polymerase chain reaction
PCD: Primary ciliary dyskinesia
RS: Radial spoke
RSP: Radial spoke protein
RS3S: Radial spoke 3 stump
MS/MS: Tandem mass spectrometry
TEM: Transmission electron microscopy
TAP: Tris-acetate-phosphate
WT: Wild-type

## Acknowledgements

We thank LeeAnn Higgins, Todd Markowski, Bruce Witthun, and Alan Zimmerman in the Center for Mass Spectrometry and Proteomics at the University of Minnesota (UMN) for expert assistance with iTRAQ labeling, mass spectrometry, and spectral counting. This center is supported by multiple grants including the National Science Foundation major research instrumentation grants 9871237 and 0215759 as described at https://cbs.umn.edu/cmsp/about. We also thank Matt Laudon and the Chlamydomonas Resource Center (https://www.chlamycollection.org/) for strains. This center is supported by the National Science Foundation Living Stock Collections for Biological Research program (NSF grants 0951671 and 00017383). The Porter laboratory also acknowledges the dedicated assistance of multiple UMN undergraduates including Aimee DeCathelineau, Jasjeet Sekhon, Jared Rieck, Shada Ahrar, and Alexandria Schauer and Clare Palmer from Wesleyan University. Expert assistance with TEM and image analysis was also provided by Amanda Bednarz, Thomas Giddings, and Dr. David Mastronarde at the Boulder Laboratory for 3D Fine Structure (University of Colorado). We also thank Chen Xu (Brandeis University) and Daniel Stoddard (UT Southwestern Medical Center) for dedicated training and maintenance of EM facilities. The UTSW cryo-electron microscope facility is funded in part by a CPRIT Core Facility Award (RP170644). Richard Linck (UMN), Toshiki Yagi (Prefectural University of Hiroshima), Win Sale (Emory University), Ritsu Kamiya (Tokyo University), Paul Lefebvre (UMN), and Pinfen Yang (Marquette University) generously supplied antibodies as listed in Supplemental Table 3. Preliminary reports of this work were presented at the American Society for Cell Biology meetings and the International Conference on the Cell and Molecular Biology of *Chlamydomonas.* This work was supported by National Institutes of Health grants to M.E.P (GM-055667), D.N. (GM-088122), and E.F.S. (GM112050). The authors declare no competing financial interests.

## Author contributions

M.E.P and D.N. designed research and obtained funding. J.L. and G.F. carried out axoneme preparation, cryoelectron tomography, and sub-tomogram averaging. T.L. cloned *FAP57* and characterized the *FAP57* gene, *ida8* mutants, rescued strains, and FAP57 antibody. T.L., C.P., K.V.M., R.B., and K.A. isolated axonemes, performed Western blot and iTRAQ analyses, carried out immunofluorescence studies, and measured swimming velocities. D.T. characterized the *bop2-1* mutation, generated all epitope-tagged constructs, and transformed and screened all mutant strains. E.O. analyzed axonemes by conventional transmission electron microscopy and 2D image averaging. E.D. and E.S. measured microtubule sliding velocities and recorded flagellar waveforms. M.E.P., J.L., and D.N. wrote the paper with input from all authors.

## Supplemental Information for

**Supplemental Figure 1.**
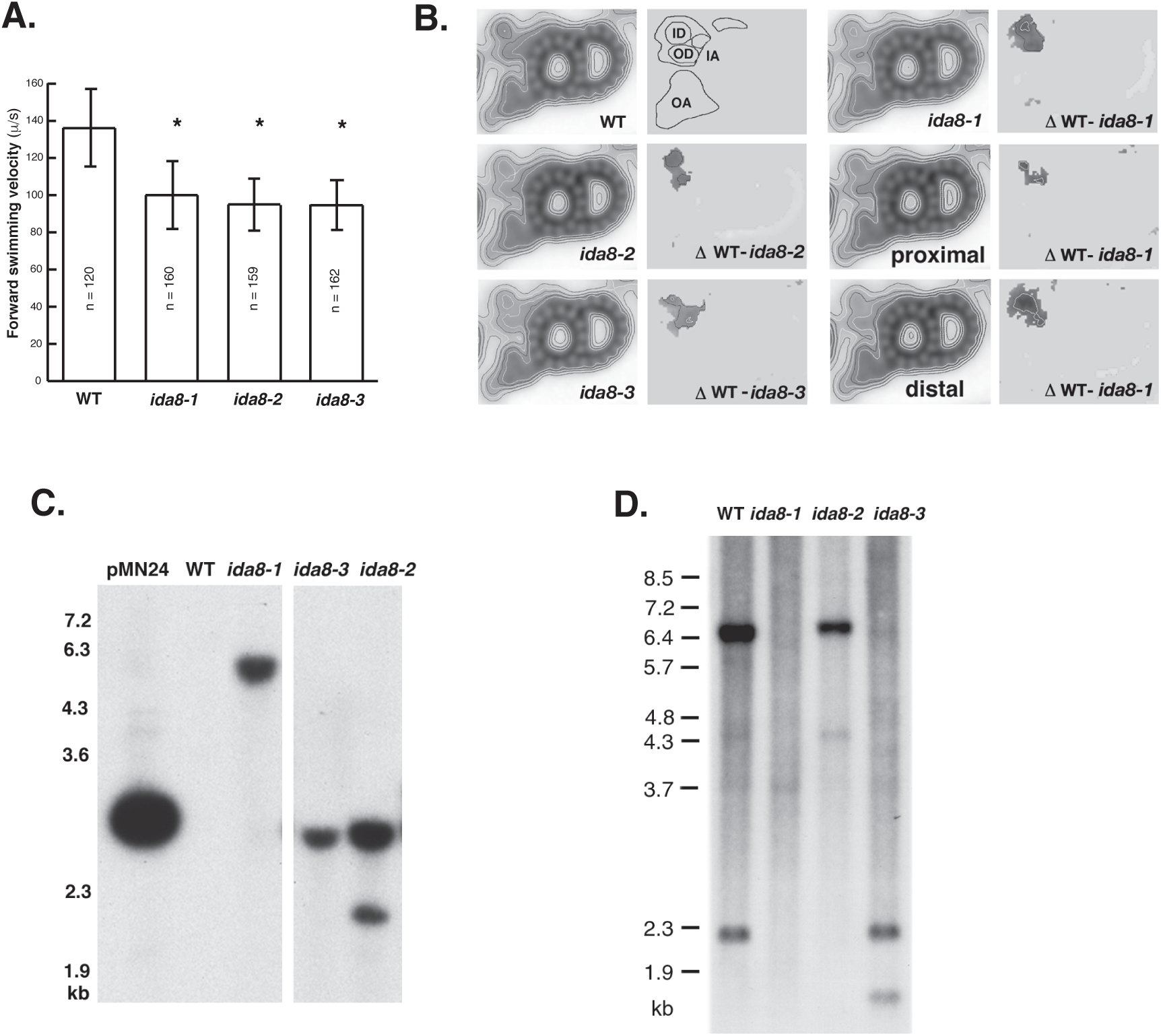
Characterization of insertional mutations in the *IDA8* locus. **(A)** The forward swimming velocities of three insertional mutant strains were measured by phase contrast light microscopy. The three strains were all significantly slower (P<0.05) than wild-type cells. **(B)** Potential defects in structure in the inner dynein arm (IDA) region were assessed by thin section electron microscopy (TEM) and computer imaging averaging (O’Toole et al., 2005). Shown here are grand averages and difference plots of the outer DMTs viewed in cross-section, with the outer arms (OA) on the bottom and the inner arms (IA) on the top. Based on contour maps, the inner arm region contains two major domains of density, an outer domain (OD) adjacent to the outer arms and an inner domain (ID) corresponding to the location of the single-headed IDAs. The three *ida8* strains showed similar structural defects in the inner domain as illustrated by the difference plots. The outer DMTs of *ida8-1* were sorted into proximal and medial/distal cross-sections and compared to WT. The defects were more prominent in the medial/distal regions of the axoneme. The number of DMT cross-sections in each average were WT (424), *ida8-1* (659), *ida8-2* (413), and *ida8-3* (183). **(C)** Genomic DNA from WT and three *ida8* strains was digested with *PvuII* and analyzed on a Southern blot probed with the pUC119 vector. **(D)** Genomic DNA from WT and three *ida8* strains was digested with *SacI* and probed with a 10 kb *NotI* fragment from the region that contains the *IDA8* gene (see Figure 1A). This fragment hybridized with 2.3 kb and 6.5 kb *SacI* restriction fragments in WT DNA. Both fragments were missing in *ida8-1*; the 2.3 kb fragment was missing in *ida8-2*, and the 6.5 kb fragment was missing in *ida8-3*. See Figure 1A for summary.

**Supplemental Figure 2.**
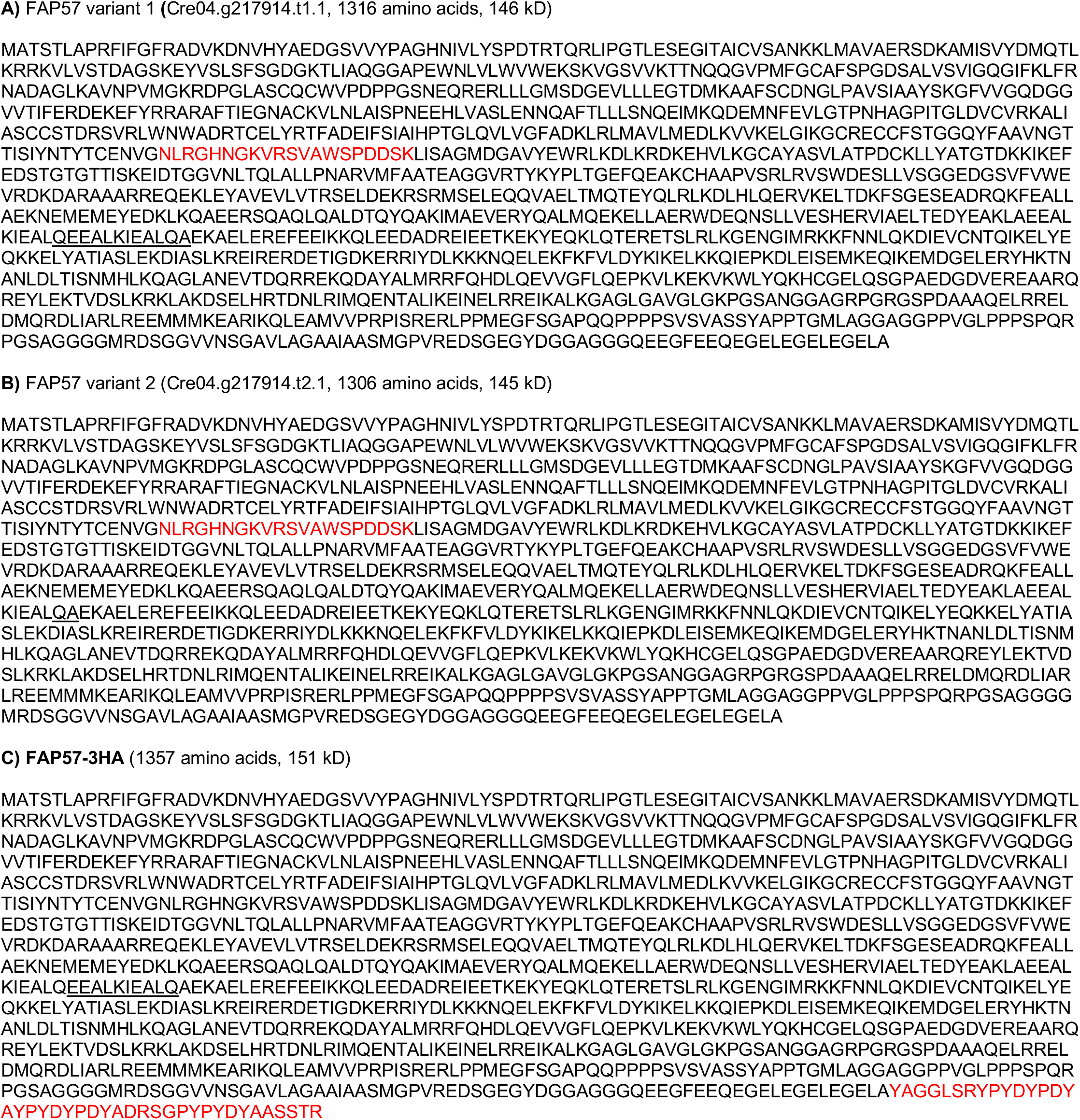
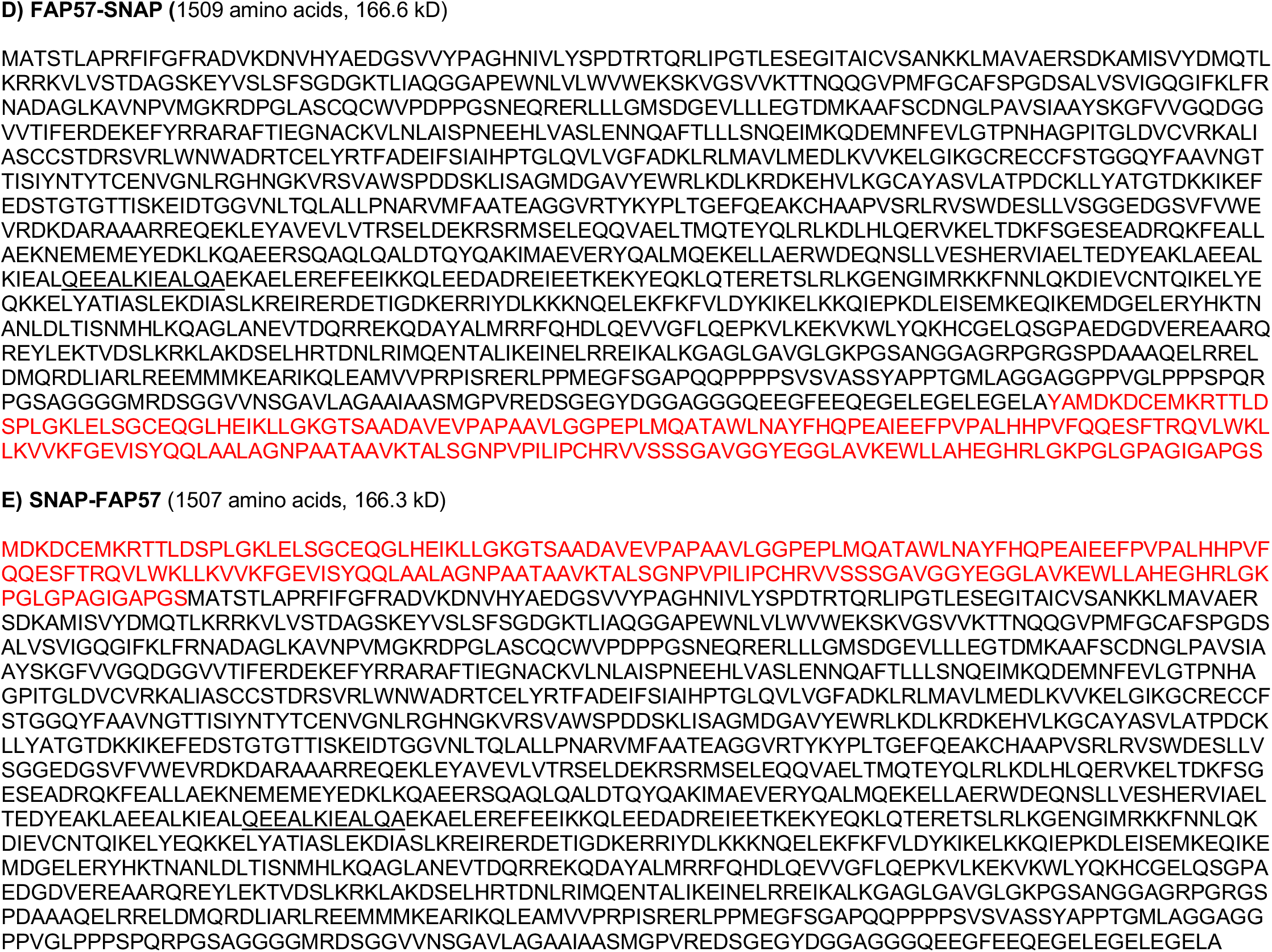
Predicted amino acid sequences of FAP57 and its tagged variants. The predicted amino acid sequences of the **(A, B)** two FAP57 polypeptides generated by alternative splicing (see underlined amino acids) are shown. Also shown are epitope-tagged versions containing **(C)** a triple HA tag at the C-terminus, **(D)** a SNAP tag at the C-terminus, or **(E)** a SNAP tag at the N-terminus. The peptide sequence used to generate a FAP57 specific antibody is shown in red in **(A)**, and the sequences of the epitope tags are shown in red in (**C, D,** and **E).**

**Supplemental figure 3.**
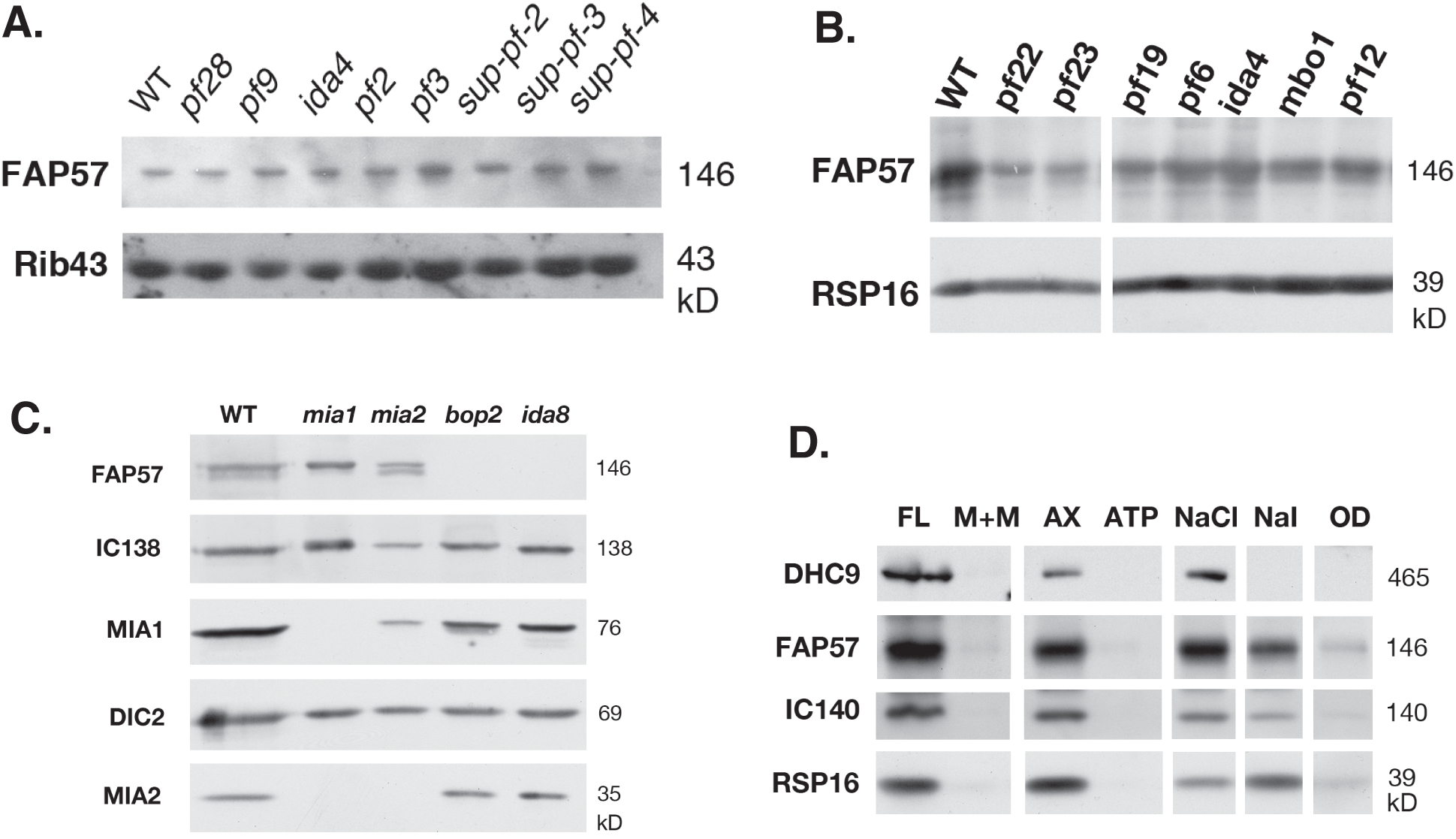
The distribution of the FAP57 polypeptide in different motility mutants and flagellar extracts. **(A)** A Western blot of axonemes from several mutants with defects in the ODAs, IDAs, or N-DRC was probed with antibodies against FAP57 and Rib43 as a loading control. **(B)** A Western blot of axonemes from several mutants with multiple dynein defects, central pair defects, or other uncharacterized motility defects was probed with antibodies against FAP57 and RSP16 as a loading control. **(C)** A Western blot of axonemes from the *mia1, mia2, bop2-1,* and *ida8-1* strains was probed with antibodies against FAP57, IC138, MIA1, DIC2 (IC69), and MIA2. **(D)** A Western blot of flagella (FL), a detergent-extracted, membrane plus matrix fraction (M+M), axonemes (AX), a 10 mM Mg ATP extract (ATP), a 0.6M NaCl extract (NaCl), a 0.5M NaI extract (NaI), and the final pellet of extracted outer doublets (OD) was probed with antibodies against DHC9, FAP57, the I1 subunit IC140, and the radial spoke subunit RSP16. Most of the FAP57 protein was extracted with 0.6M NaCl and the remainder with 0.6M NaI.

**Supplemental figure 4.**
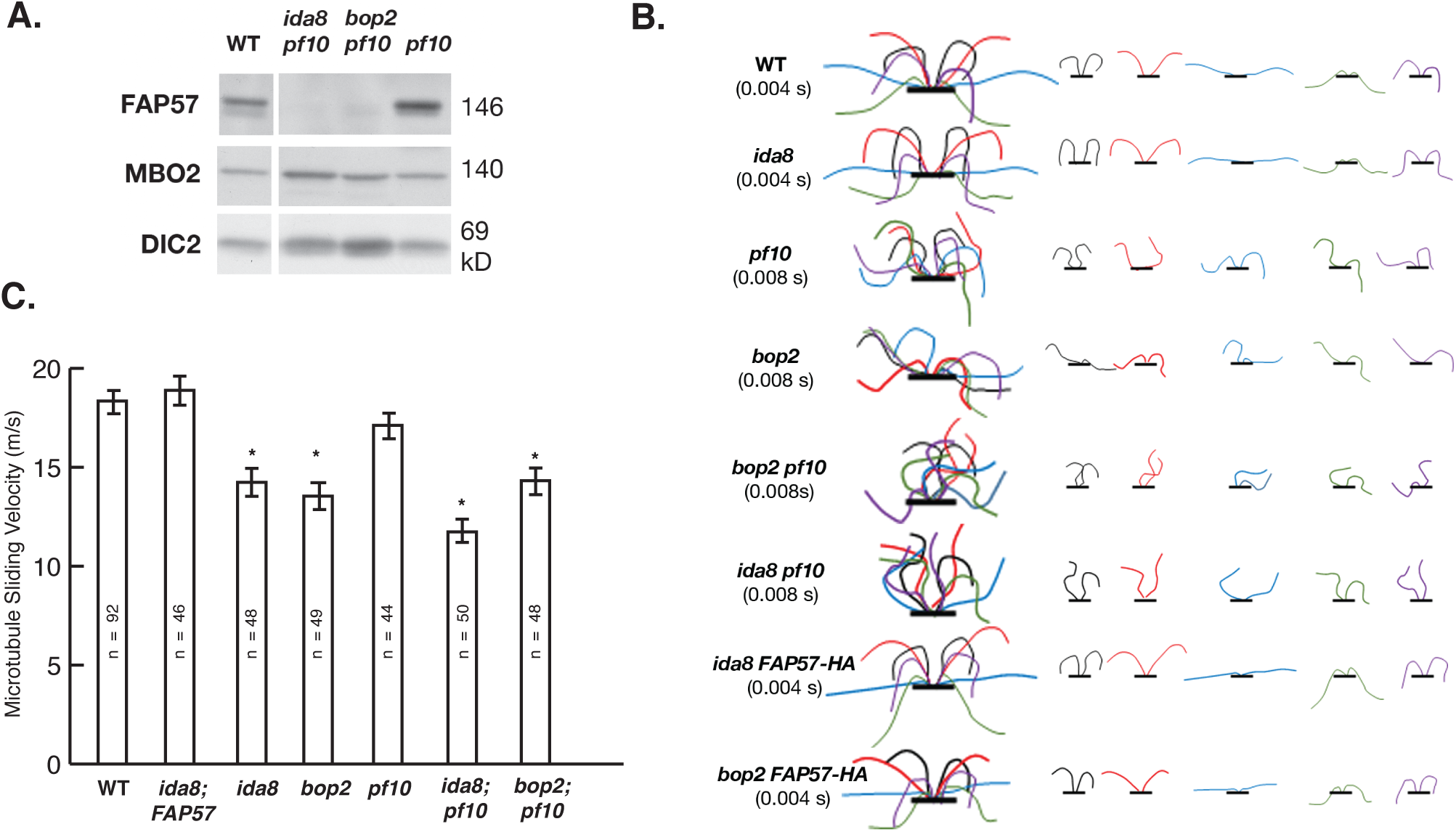
Phenotypic interactions between *pf10, bop2-1,* and *ida8-1*. **(A)** A Western blot of axonemes from wild-type, *pf10*, and the double mutants *ida8-1; pf10* and *bop2-1; pf10* was probed with antibodies against FAP57, MBO2, and DIC2 (IC69) as a loading control. **(B)** Tracings of the flagellar waveforms observed in high speed videos of WT, *ida8-1, pf10, bop2-1, pf10* double mutants, and *FAP57-HA* rescued strains are shown here. The original videos are shown in Supplemental Videos 1-8. **(C)** Measurements of microtubule sliding velocities observed during protease-induced sliding disintegration of isolated axonemes from different strains. Values shown are mean +/- SEM. The sliding velocities of *ida8* and *bop2* were significantly slower (P < 0.05) than those of the WT, the FAP57 rescued strain, and *pf10*, but not significantly different from one another. The sliding velocities of *ida8; pf10* and *bop2; pf10* were also significantly slower (P < 0.05) than that of *pf10* alone.

**Supplemental Figure 5.**
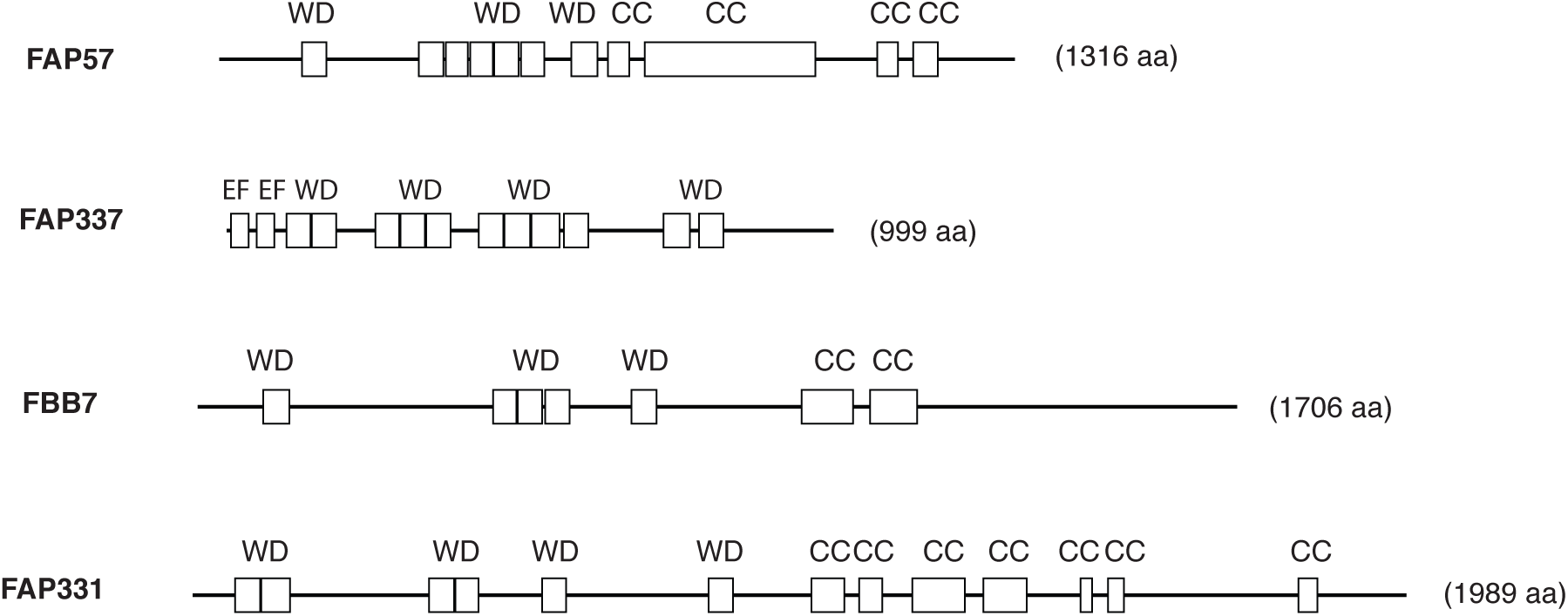
Diagrams of polypeptides that are altered in *ida8* axonemes as determined by iTRAQ labeling and tandem MS/MS. The relative sizes of the polypeptides and their predicted structural domains are drawn to scale. WD repeat domains (WD), coiled-coil domains (CC), and EF hand domains (EF).

**Supplemental Figure 6.**
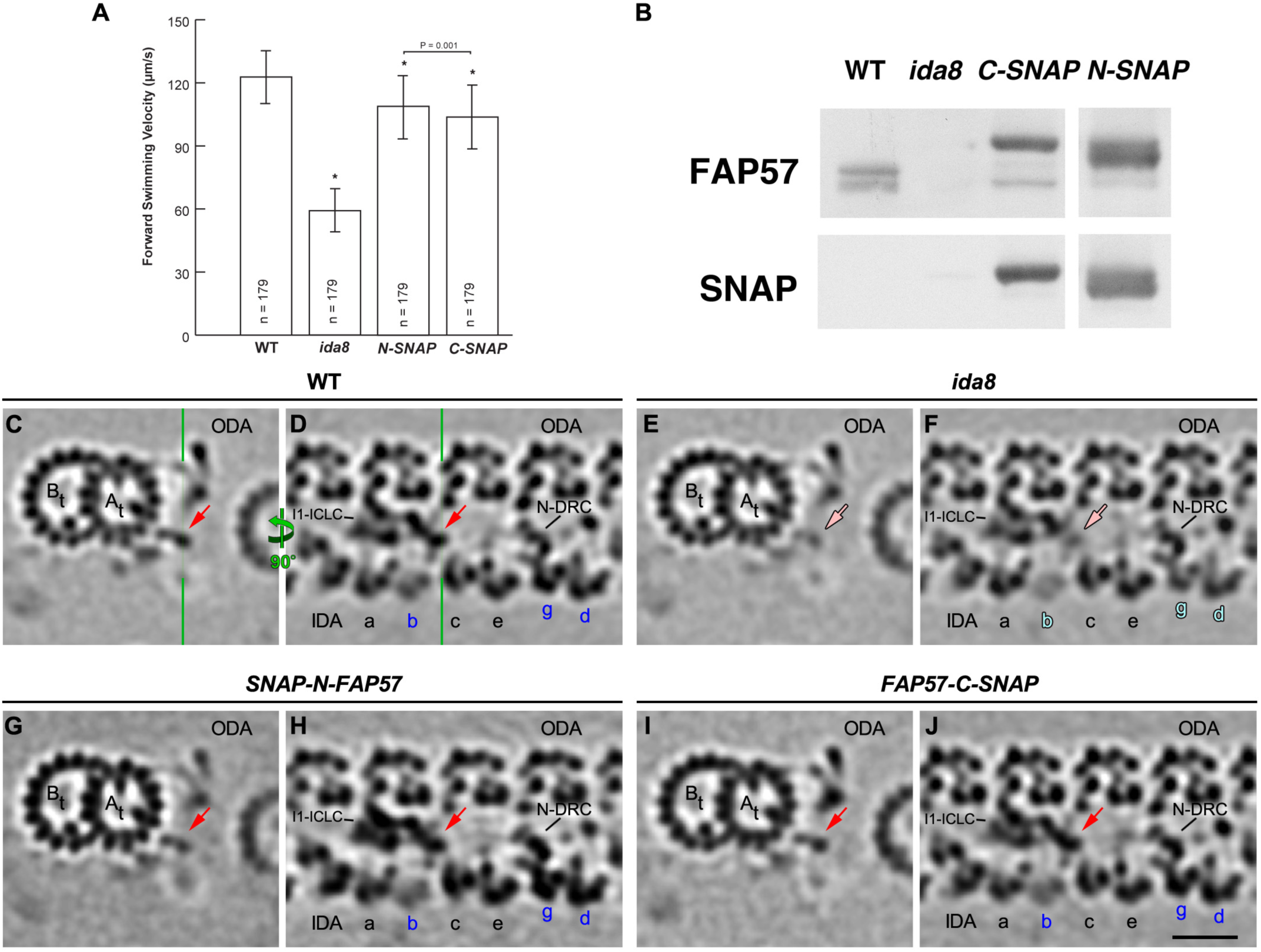
Rescue of *ida8* with SNAP-tagged FAP57 constructs. **(A)** Measurements of forward swimming velocities by phase contrast microscopy. FAP57 constructs containing either an N-terminal SNAP tag (*N-SNAP*) or C-terminal SNAP tag (*C-SNAP*) increased the forward swimming velocities of *ida8-1* rescued strains to near wild-type levels. **(B)** A Western blot of axonemes from WT, *ida8,* and the *C-SNAP* and *N-SNAP FAP57* rescued strains was probed with antibodies against FAP57 or the SNAP tag. Both constructs were assembled efficiently into axonemes. **(C-J**) Comparison of the averaged 96 nm repeats revealed the structural defects in *ida8* and the rescue of *ida8* with SNAP-tagged FAP57 constructs. Tomographic slices were taken from cross sections of the 96 nm repeat at the position of the I1-distal structure (C, E, G, I) or in longitudinal sections through the dyneins with the ODAs on the top and the IDAs at the bottom (D, F, H, J). The arrows indicate the I1-distal structure present in WT (C, D; red), reduced in *ida8*, (E, F; pink), and recovered in the *SNAP-N-FAP57* rescued strain (G, H; red), and the *FAP57-C-SNAP* rescued strain (I, J; red). Analysis of all *ida8* tomograms also suggested defects in the assembly of IDAs *b, g*, and *d*. However, more detailed classification analyses only confirmed the defects of I1-distal structure and IDAs *g* and *d* in *ida8,* and their recovery in rescue strains. Scale bar in (J) is 20 nm.

**Supplemental Figure 7.**
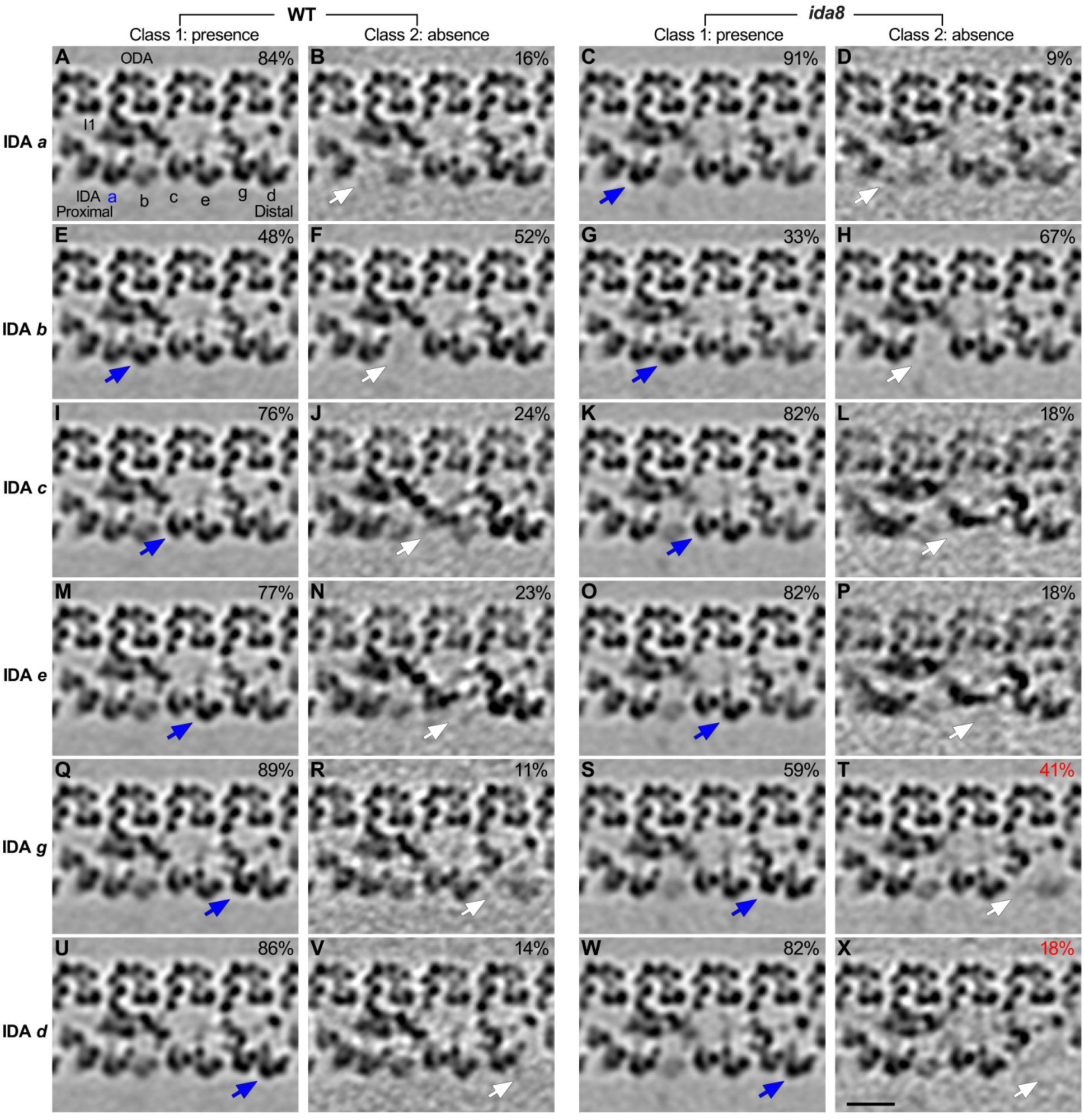
Comparison of class averages of the 96 nm repeats from WT and *ida8* reveals defects in the assembly of IDAs *d* and *g* in *ida8* axonemes. Classification analysis was performed on the 96 nm repeats from WT and *ida8* axonemes for the presence (Class 1) or absence (Class 2) of each single-headed IDA (*a, b, c, e, g, d*). Longitudinal tomographic slices of the 96 nm repeats with the four ODAs on top and the six single-headed IDAs at the bottom are shown here. The presence and absence of a given IDA structure that is indicated on the left are highlighted by blue and white arrows, respectively. The percentage of sub-tomograms included in each class average is also indicated. Note that although the percentage of tomograms that lack IDA *b* was increased in *ida8* (F, H), this increase was primarily due to the higher proportion of tomograms from the proximal region in the *ida8* dataset (see Figure 7 and Discussion). Scale bar in (X) is 20 nm.

**Supplemental Figure 8.**
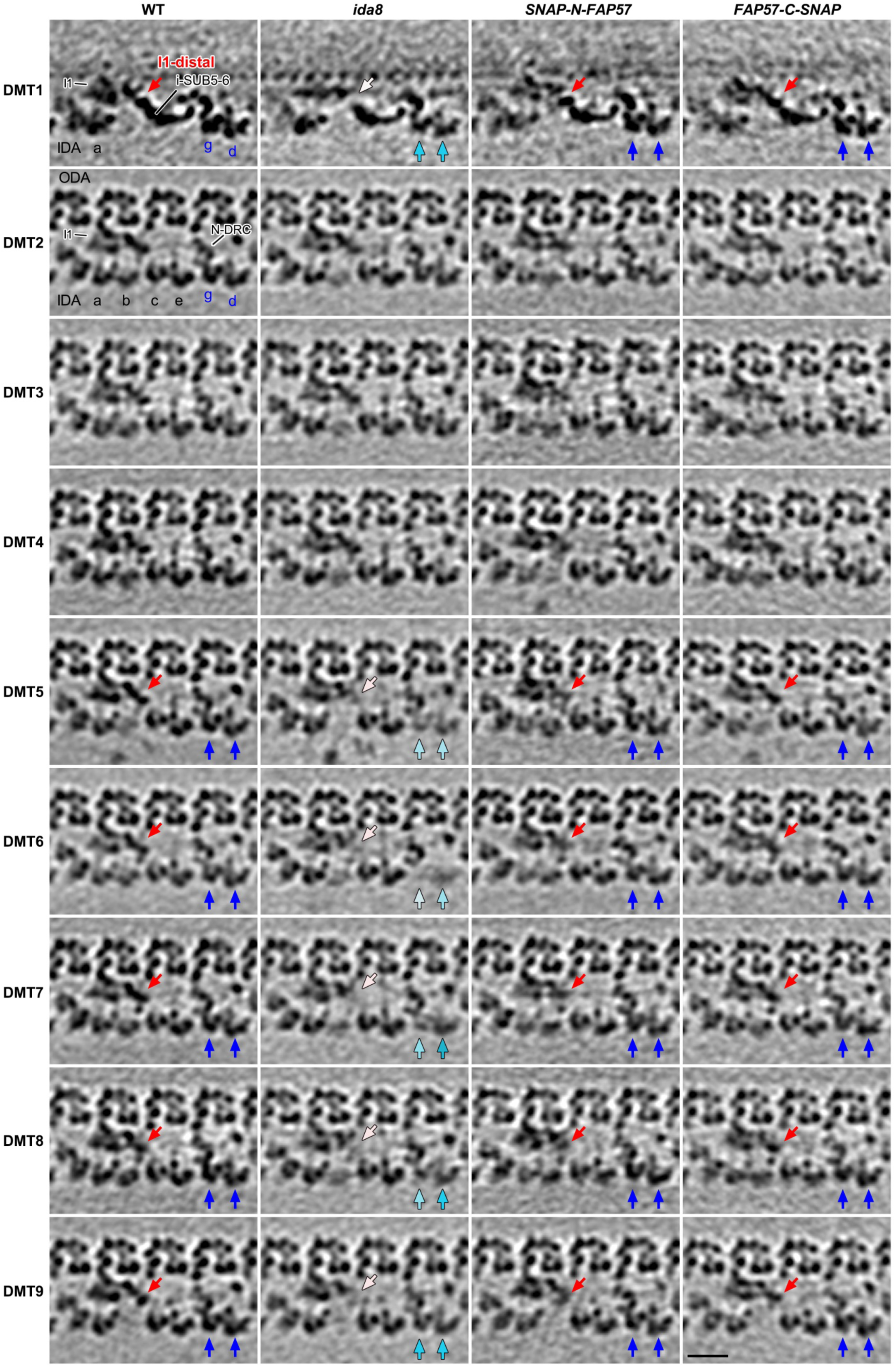
DMT specific averaging reveals the asymmetric distribution of structural defects in *ida8* and their recovery in SNAP-tagged *FAP57* strains. The 96 nm repeats on each individual DMT (1-9) were averaged for each strain (WT, *ida8, SNAP-N-FAP57* and *FAP57-C-SNAP*). Shown here are tomographic slices that were taken from longitudinal sections of the 96 nm repeats, with the ODAs on the top and the IDAs at the bottom. The red arrows indicate the normal I1-distal structure in WT and its recovery in the two rescued strains, *SNAP-N-FAP57* and *FAP57-C-SNAP*. The lighter pink arrows highlight the reduction of the I1-distal structure on DMTs 1, 5-9 in *ida8*. The dark blue arrows indicate the normal assembly of IDAs *g* and *d* in WT and the two rescued strains, *SNAP-N-FAP57* and *FAP57-C-SNAP*. The lighter blue arrows highlight the obvious reduction in the densities of IDAs *g* and *d* on DMTs 1, 5-9 in *ida8*. Scale bar is 20 nm.

**Supplemental Table 1.**
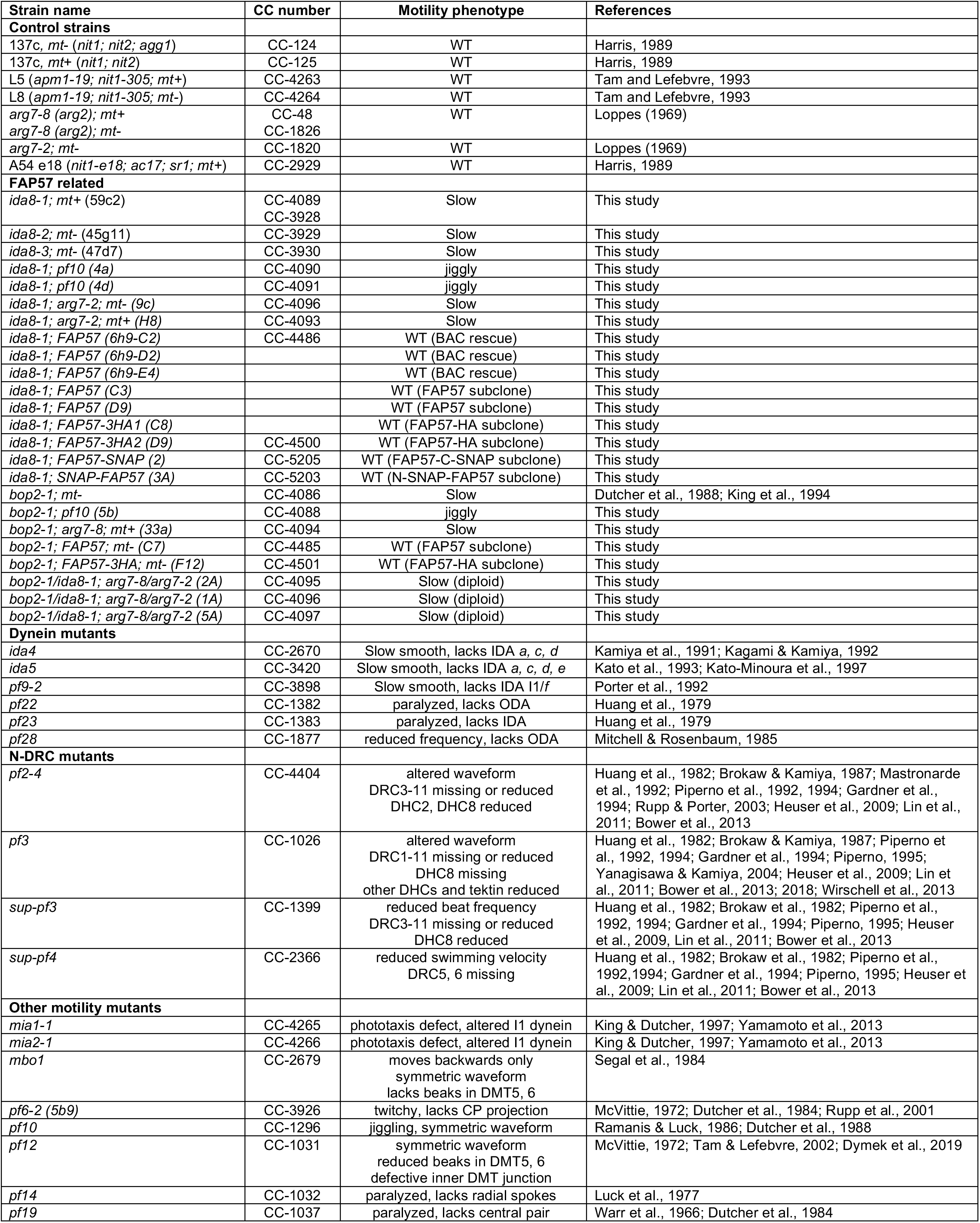
Strains used in this study.

| Strain name | CC number | Motility phenotype | References |
| --- | --- | --- | --- |
| **Control strains** |  |  |  |
| 137c, *mt-* (*nit1*; *nit2*; *agg1*) | CC-124 | WT | Harris, 1989 |
| 137c, *mt+* (*nit1*; *nit2*) | CC-125 | WT | Harris, 1989 |
| L5 (*apm1-19*; *nit1-305*; *mt+*) | CC-4263 | WT | Tam and Lefebvre, 1993 |
| L8 (*apm1-19*; *nit1-305*; *mt-*) | CC-4264 | WT | Tam and Lefebvre, 1993 |
| *arg7-8* (*arg2*); *mt+* | CC-48 | WT | Loppes (1969) |
| *arg7-8* (*arg2*); *mt-* | CC-1826 |  |  |
| *arg7-2*; *mt-* | CC-1820 | WT | Loppes (1969) |
| A54 e18 (*nit1-e18*; *ac17*; *sr1*; *mt+*) | CC-2929 | WT | Harris, 1989 |
| **FAP57 related** |  |  |  |
| *ida8-1*; *mt+* (59c2) | CC-4089<br>CC-3928 | Slow | This study |
| *ida8-2*; *mt-* (45g11) | CC-3929 | Slow | This study |
| *ida8-3*; *mt-* (47d7) | CC-3930 | Slow | This study |
| *ida8-1*; *pf10* (4a) | CC-4090 | jiggly | This study |
| *ida8-1*; *pf10* (4d) | CC-4091 | jiggly | This study |
| *ida8-1*; *arg7-2*; *mt-* (9c) | CC-4096 | Slow | This study |
| *ida8-1*; *arg7-2*; *mt+* (H8) | CC-4093 | Slow | This study |
| *ida8-1*; FAP57 (6h9-C2) | CC-4486 | WT (BAC rescue) | This study |
| *ida8-1*; FAP57 (6h9-D2) |  | WT (BAC rescue) | This study |
| *ida8-1*; FAP57 (6h9-E4) |  | WT (BAC rescue) | This study |
| *ida8-1*; FAP57 (C3) |  | WT (FAP57 subclone) | This study |
| *ida8-1*; FAP57 (D9) |  | WT (FAP57 subclone) | This study |
| *ida8-1*; FAP57-3HA1 (C8) |  | WT (FAP57-HA subclone) | This study |
| *ida8-1*; FAP57-3HA2 (D9) | CC-4500 | WT (FAP57-HA subclone) | This study |
| *ida8-1*; FAP57-SNAP (2) | CC-5205 | WT (FAP57-C-SNAP subclone) | This study |
| *ida8-1*; SNAP-FAP57 (3A) | CC-5203 | WT (N-SNAP-FAP57 subclone) | This study |
| *bop2-1*; *mt-* | CC-4086 | Slow | Dutcher et al., 1988; King et al., 1994 |
| *bop2-1*; *pf10* (5b) | CC-4088 | jiggly | This study |
| *bop2-1*; *arg7-8*; *mt+* (33a) | CC-4094 | Slow | This study |
| *bop2-1*; FAP57; *mt-* (C7) | CC-4485 | WT (FAP57 subclone) | This study |
| *bop2-1*; FAP57-3HA; *mt-* (F12) | CC-4501 | WT (FAP57-HA subclone) | This study |
| *bop2-1/ida8-1*; *arg7-8/arg7-2* (2A) | CC-4095 | Slow (diploid) | This study |
| *bop2-1/ida8-1*; *arg7-8/arg7-2* (1A) | CC-4096 | Slow (diploid) | This study |
| *bop2-1/ida8-1*; *arg7-8/arg7-2* (5A) | CC-4097 | Slow (diploid) | This study |
| **Dynein mutants** |  |  |  |
| *ida4* | CC-2670 | Slow smooth, lacks IDA *a*, *c*, *d* | Kamiya et al., 1991; Kagami & Kamiya, 1992 |
| *ida5* | CC-3420 | Slow smooth, lacks IDA *a*, *c*, *d*, *e* | Kato et al., 1993; Kato-Minoura et al., 1997 |
| *pf9-2* | CC-3898 | Slow smooth, lacks IDA I1/f | Porter et al., 1992 |
| *pf22* | CC-1382 | paralyzed, lacks ODA | Huang et al., 1979 |
| *pf23* | CC-1383 | paralyzed, lacks IDA | Huang et al., 1979 |
| *pf28* | CC-1877 | reduced frequency, lacks ODA | Mitchell & Rosenbaum, 1985 |
| **N-DRC mutants** |  |  |  |
| *pf2-4* | CC-4404 | altered waveform<br>DRC3-11 missing or reduced<br>DHC2, DHC8 reduced | Huang et al., 1982; Brokaw & Kamiya, 1987; Mastronarde et al., 1992; Piperno et al., 1992, 1994; Gardner et al., 1994; Rupp & Porter, 2003; Heuser et al., 2009; Lin et al., 2011; Bower et al., 2013 |
| *pf3* | CC-1026 | altered waveform<br>DRC1-11 missing or reduced<br>DHC8 missing<br>other DHCs and tektin reduced | Huang et al., 1982; Brokaw & Kamiya, 1987; Piperno et al., 1992, 1994; Gardner et al., 1994; Piperno, 1995; Yanagisawa & Kamiya, 2004; Heuser et al., 2009; Lin et al., 2011; Bower et al., 2013; 2018; Wirschell et al., 2013 |
| *sup-pf3* | CC-1399 | reduced beat frequency<br>DRC3-11 missing or reduced<br>DHC8 reduced | Huang et al., 1982; Brokaw et al., 1982; Piperno et al., 1992, 1994; Gardner et al., 1994; Piperno, 1995; Heuser et al., 2009; Lin et al., 2011; Bower et al., 2013 |
| *sup-pf4* | CC-2366 | reduced swimming velocity<br>DRC5, 6 missing | Huang et al., 1982; Brokaw et al., 1982; Piperno et al., 1992, 1994; Gardner et al., 1994; Piperno, 1995; Heuser et al., 2009; Lin et al., 2011; Bower et al., 2013 |
| **Other motility mutants** |  |  |  |
| *mia1-1* | CC-4265 | phototaxis defect, altered I1 dynein | King & Dutcher, 1997; Yamamoto et al., 2013 |
| *mia2-1* | CC-4266 | phototaxis defect, altered I1 dynein | King & Dutcher, 1997; Yamamoto et al., 2013 |
| *mbo1* | CC-2679 | moves backwards only<br>symmetric waveform<br>lacks beaks in DMT5, 6 | Segal et al., 1984 |
| *pf6-2* (5b9) | CC-3926 | twitchy, lacks CP projection | McVittie, 1972; Dutcher et al., 1984; Rupp et al., 2001 |
| *pf10* | CC-1296 | jiggling, symmetric waveform | Ramanis & Luck, 1986; Dutcher et al., 1988 |
| *pf12* | CC-1031 | symmetric waveform<br>reduced beaks in DMT5, 6<br>defective inner DMT junction | McVittie, 1972; Tam & Lefebvre, 2002; Dymek et al., 2019 |
| *pf14* | CC-1032 | paralyzed, lacks radial spokes | Luck et al., 1977 |
| *pf19* | CC-1037 | paralyzed, lacks central pair | Warr et al., 1966; Dutcher et al., 1984 |

**Supplemental Table 2.**
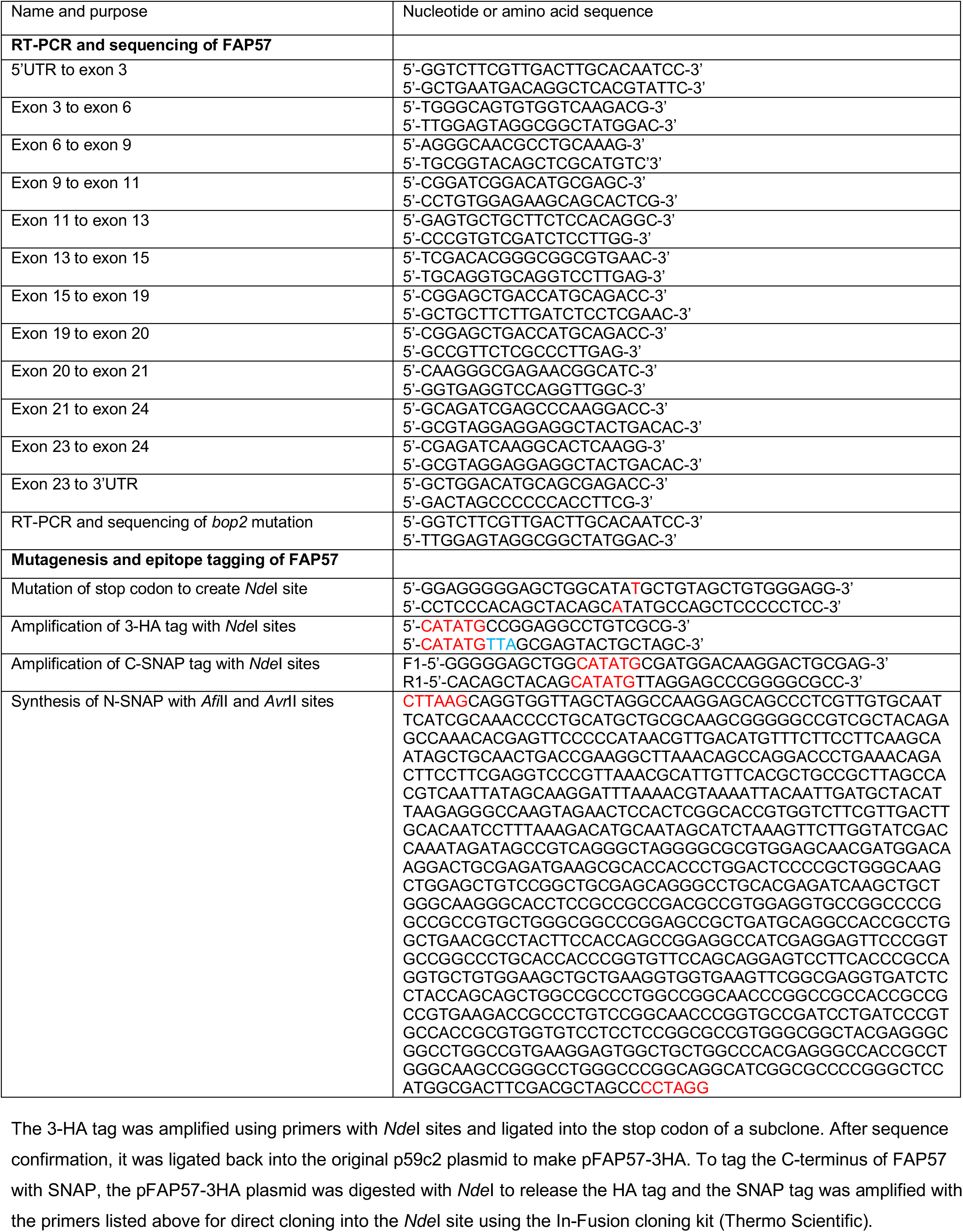
Oligonucleotide sequences used in the study.

**Supplemental Table 3.**
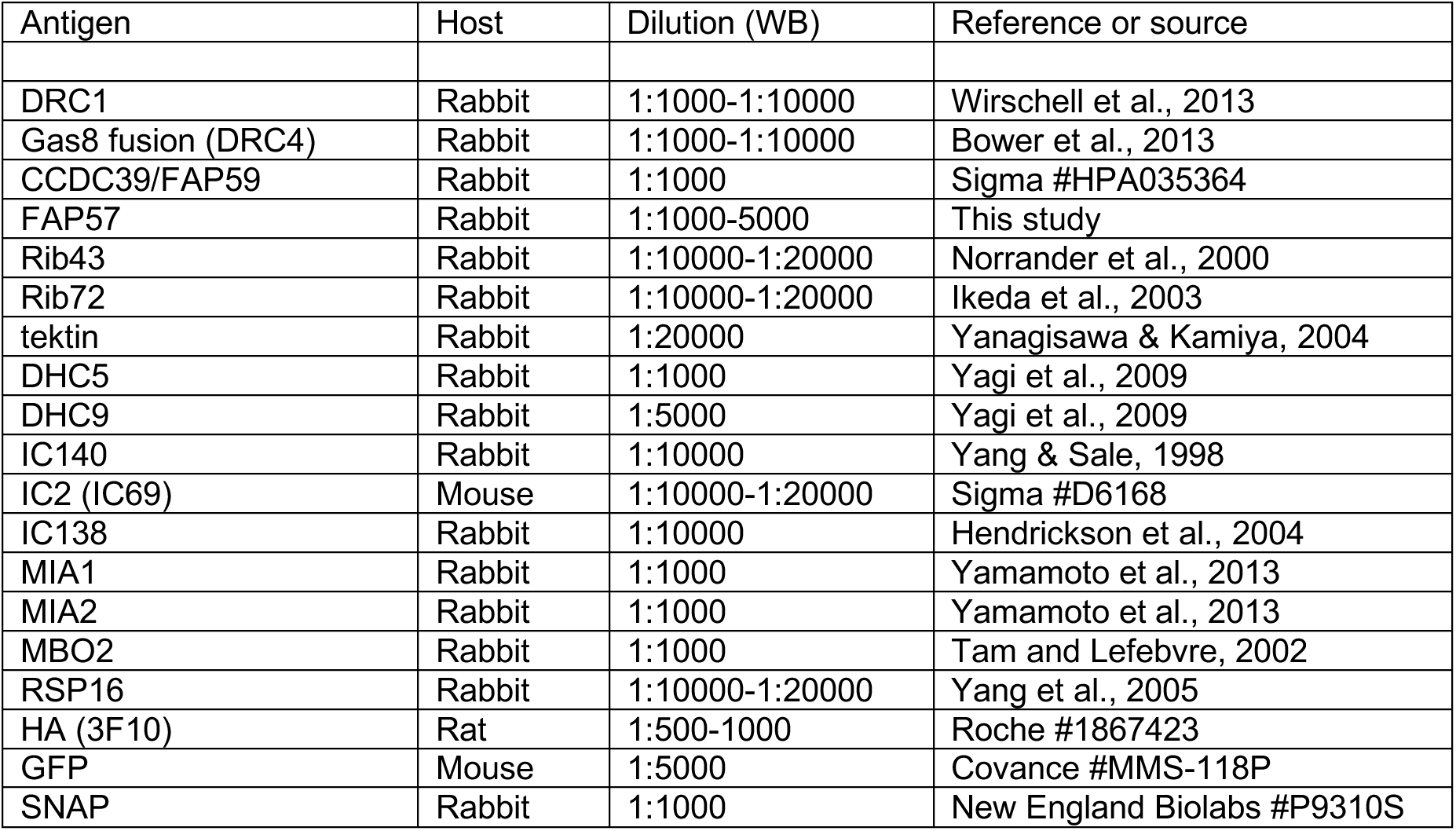
Antibodies used in this study.

**Supplemental Table 4.** Polypeptides co-eluting with FAP57 in FPLC peak *g*.

| **Protein** | ***Chlamydomonas* ID** | **Peptides (N)** | **Length (aa) and MW (kD)** | **Predicted domains** | **Best hit in *H. sapiens*<sup>a</sup>** |
| --- | --- | --- | --- | --- | --- |
| FAP44 | Cre09.g386736 | 44 | 2141 (222) | WD, CC | WDR52 |
| FAP43 | Cre16.g691440 | 34 | 1950 (199) | WD, CC | WDR96 |
| FAP57 | Cre04.g217914 | 24 | 1316 (146) | WD, CC | WDR65 |
| DHC7 | Cre14.g627576 | 32 | 4191 (468) | AAA, CC | DHC9A |
| FAP244 | Cre08.g374700 | 7 | 2630 (267) | WD, CC | N.D. |
| IFT88 | Cre07.g335750 | 7 | 782 (86) | TPR | IFT88 |
| 1α DHC | Cre12.g484250 | 5 | 4626 (523) | AAA, CC | DHC5 |
| Actin | Cre13.g603700 | 5 | 377 (42) |  | Actin |
| FAP159 | Cre01.g022750 | 3 | 2577 (252) | LRR | N.D. |
| FAP75 | Cre06.g249900 | 3 | 1264 (125) | AAA | AK7 |
| IC97 (DII6) | Cre14.g631200 | 3 | 760 (82) | CC | CASC1 |

<sup>a</sup>DHCs in *H. sapiens* are named according to their DHC class as defined by Kollmar, 2000.

**Table S5.** *Chlamydomonas* strains used for cryo ET analyses.

| Strain | Number of tomograms | Averaged repeats | Resolution 0.5 FSC (nm) |
| --- | --- | --- | --- |
| WT-I<sup>a</sup>) (F30/CCD data) | 25 | 3736 | 3.4 |
| WT-II<sup>b</sup>) (Krios/K2/VPP data; higher resolution) | 80 | 11225 | 1.8 |
| *ida8* | 33 | 4227 | 3.8 |
| *SNAP-N-FAP57* control | 12 | 1500 | 3.6 |
| *SNAP-N-FAP57*+Au | 18 | 2207 | 3.6 |
| *FAP57-C-SNAP* control | 10 | 1500 | 4.1 |
| *FAP57-C-SNAP*+Au | 16 | 2498 | 3.7 |

a) The WT-I reference dataset is a composite of tomograms from WT (CC-125), *ida6::IDA6-GFP* (CC-4495), *DRC3-SNAP* (CC-5243), and *SNAP-DRC3* (CC-5244) axonemes. Some of these tomograms were previously used to analyze other axonemal complexes (Song et al., 2015; Bower et al., 2018). The *ida6::IDA6-GFP* strain was generated by rescuing *ida6* with a GFP-tagged WT *IDA6* gene (Bower et al., 2018). The *DRC3-SNAP* and *SNAP-DRC3* strains were generated by rescuing *drc3* with the WT *DRC3* gene tagged with SNAP at either its N-terminal or C-terminal end (Awata et al., 2015; Song et al., 2015). All of these axonemes are structurally and phenotypically indistinguishable from WT.
b) The WT-II reference dataset is a composite of tomograms from WT (CC-4533) and several central pair mutants (*fap76-1*, *fap81*, *fap92*, *fap216*, and *fap76-1; fap81*) obtained from the *Chlamydomonas* CLiP library (CLiP ID numbers: LMJ.RY0402.089534, LMJ.RY0402.092632, LMJ.RY0402.204383, and LMJ.RY0402.218389, respectively). Some of these tomograms were previously used to analyze other axonemal complexes (Fu et al. 2019). The 96nm axoneme repeats in the central pair mutants are structurally indistinguishable from WT.

### Captions for Videos S1 to S8

Supplemental Video S1. Video of a wild-type cell swimming forward with an asymmetric waveform.

Supplemental Video S2. Video of an *ida8-1* cell swimming forward with an asymmetric waveform.

Supplemental Video S3. Video of a *bop2-1* cell swimming forward with an asymmetric waveform.

Supplemental Video S4. Video of an *ida8-1; Fap57-HA* rescued cell swimming forward with an asymmetric waveform.

Supplemental Video S5. Video of a *bop2-1, FAP57-HA* rescued cell swimming forward with an asymmetric waveform.

Supplemental Video S6. Video of a *pf10* cell swimming with an abnormal waveform.

Supplemental Video S7. Video of an *ida8-1; pf10* cell swimming with a variable waveform.

Supplemental Video S8. Video of a *bop2-1; pf10* cell swimming with a variable waveform.

